# GZMK^high^ CD8^+^ T effector memory cells are associated with CD15^high^ neutrophil abundance in early-stage colorectal tumors and predict poor clinical outcome

**DOI:** 10.1101/2021.12.14.472046

**Authors:** Silvia Tiberti, Carlotta Catozzi, Caterina Scirgolea, Ottavio Croci, Mattia Ballerini, Danilo Cagnina, Chiara Soriani, Carina B. Nava Lauson, Angeli D. Macandog, Giovanni Bertalot, Wanda L. Petz, Simona P. Ravenda, Valerio Licursi, Paola Paci, Marco Rasponi, Nicola Fazio, Guangwen Ren, Uberto Fumagalli-Romario, Martin H. Shaefer, Stefano Campaner, Enrico Lugli, Luigi Nezi, Teresa Manzo

## Abstract

Tumor contexture has emerged as a major prognostic determinant and tumor infiltrating CD8^+^ T cells have been associated with a better prognosis in several solid tumors, including early-stage colorectal cancer (CRC). However, the tumor immune infiltrate is highly heterogeneous and understanding how the interplay between different immune cell compartments impacts on the clinical outcome is still in its infancy.

Here, we describe in a prospective cohort a novel CD8^+^ T effector memory population, which is characterized by high levels of Granzyme K (GZMK^high^ CD8^+^ T_EM_) and is correlated with CD15^high^ tumor infiltrating neutrophils. We provide both *in vitro* and *in vivo* evidence of the role of stromal cell-derived factor 1 (CXCL12/SDF-1) in driving functional changes on neutrophils at the tumor site, promoting their retention and increasing the crosstalk with CD8^+^ T cells. Mechanistically, as a consequence of the interaction with neutrophils, CD8^+^ T cells are skewed towards a CD8^+^ T_EM_ phenotype and produce high levels of GZMK, which in turn decreases E-cadherin pathway. The correlations of GZMK^high^ CD8^+^ T_EM_ and neutrophils with both tumor progression in mice and early relapse in CRC patients demonstrate the role of GZMK^high^ CD8^+^ T_EM_ in promoting malignancy. Indeed, a gene signature defining GZMK^high^ CD8^+^ T_EM_ was associated with worse prognosis on a larger independent cohort of CRC patients and a similar analysis was extended to lung cancer (TCGA).

Overall, our results highlight the emergence of GZMK^high^ CD8^+^ T_EM_ in early-stage CRC tumors as a hallmark driven by the interaction with neutrophils, which could implement current patient stratification and be targeted by novel therapeutics.

## BACKGROUND

Colorectal cancer (CRC) is the third most commonly diagnosed tumor worldwide and a leading cause of death for cancer, with the efficacy of treatment and subsequent survival largely dictated by the tumor-node-metastasis (TNM) stage at diagnosis. Tumor contexture has emerged as a major determinant to establish prognosis and guide novel therapies (Galon et al., 2014; Pagès et al., 2009a). However, despite favorable TNM staging and significant lymphocyte infiltration (Ogino et al., 2009), 30-40% of patients treated for early-stage CRC relapse and ultimately die from metastatic disease (American Cancer Society, 2018), demonstrating the inadequacy of current stratification.

In this regard, tumor infiltration by cytotoxic CD8^+^ T cells has been associated with a better prognosis in several solid tumors, including CRC (Fountzilas et al., 2018; Koch et al., 2006) and high levels of memory CD8^+^ T cells prevent early metastatic invasion and are associated with better survival (Pagès et al., 2009b), supporting the idea that the intratumor immune landscape contributes to tumor control and prevention of recurrence. However, the tumor immune infiltrate is highly heterogeneous and, although the influence of various myeloid cell populations on T cell activity has been reported (Binnewies et al., 2018; Lavin et al., 2017; Salmon et al., 2016), our understanding of the interplay between different cell compartments at the tumor site and their impact on the clinical outcome is still in its infancy. In particular, tumor-associated neutrophils represent a significant fraction of the inflammatory cells in the tumor microenvironment (TME) of many types of cancers, including CRC (Berry et al., 2017; Carus et al., 2013; Ilie et al., 2014; Li et al., 2018; Rao et al., 2012; Silvestre-Roig et al., 2019). The concept of neutrophil phenotypic and functional diversity has emerged from murine tumor models (Ballesteros et al., 2020), but the existence of subsets of infiltrating neutrophils with different functions in humans is still debated. As a result, the contribution of these cells to inhibiting or promoting tumor progression by reshaping anti-tumor immune response remains largely unexplored and often contradictory in humans (Gentles et al., 2015; Singhal et al., 2016).

Here, we sought to gain insight into the crosstalk between the TME and immune cells to elucidate their contribution to relapse and implement patient’s stratification for subsequent immune-based therapeutics. We have estimated heterogeneity of immune cells within and across CRC patients by employing multiparametric flow cytometry and single-cell transcriptomics, obtaining functional information. This combined approach has revealed an association of neutrophils infiltrating the tumor with a unique immune signature of tumor infiltrating lymphocytes (TILs), which is characterized by high levels of Granzyme K (GZMK) expression and leads to worse prognosis.

## METHODS

### Study approval

Tumors, adjacent normal tissue and peripheral blood samples from patients diagnosed with CRC at European Institute of Oncology (IEO), were collected with written informed consent as approved by the Internal Research Center Review Board. All patients were treatment-naïve at the time of sample collection (Table 1) and fresh tissue was collected the day of the surgery. All donors provided their consent in accordance with the Declaration of Helsinki. Mice were obtained from Charles River, housed and bred in a specific pathogen free animal facility. The experiments described in this manuscript were performed in accordance with the European Union Guideline on Animal Experiments. Mouse protocols were approved by the Italian Ministry of Health, the IEO Committee and the Institutional Animal Care and Use Committee of The Jackson Laboratory. Mice were matched for age and sex in each experiment.

**Table 1.** CRC patients’ characteristics. Age, Gender, Stage at first diagnosis, Body Mass Index (BMI), Site of primary tumor and Tumor type are shown for the study cohort.

| Study Cohort |  |  |
| --- | --- | --- |
|  | Count | Frequency (%) |
| Total of patients | 42 | 100 |
| Relapse (2 years) | 4 | 9,5 |
| MSI | 6 | 14,3 |
| Age |  |  |
| <50 | 2 | 4,8 |
| 50-70 | 22 | 52,4 |
| >70 | 18 | 42,9 |
| Gender |  |  |
| F | 20 | 47,6 |
| M | 22 | 52,4 |
| Stage first diagnosis |  |  |
| I | 9 | 21,4 |
| II | 15 | 35,7 |
| III | 18 | 42,9 |
| BMI |  |  |
| Normal | 19 | 45,2 |
| Overweight | 15 | 35,7 |
| Obese | 7 | 16,7 |
| NA | 1 (NT14) | 2,4 |
| Site of primary tumor |  |  |
| Left | 18 | 42,8 |
| Right | 23 | 54,8 |
| Medium Transverse Colon | 1 | 2,4 |
| Tumor Type |  |  |
| Adenocarcinoma | 39 | 92,9 |
| Mucinous adenocarcinoma | 3 | 7,1 |

### Tissue dissociation and cells

Primary tumors (T) and Normal Adjacent Tissue (NAT, collected at 10cm from the tumor site) were dissociated into single-cell suspension by mechanical and enzymatic disaggregation. The samples were then filtered through 70-μm cell strainers and washed in PBS without Ca^2+^ and Mg^2+^. Interstitial fluid was prepared from T and NAT. Briefly, tissues were minced in cold PBS and incubated on ice until cellular debris was precipitated. The supernatant was collected and spinned at 1200g at 4**°**C and then used fresh or stored at - 80**°**C.

Peripheral blood mononuclear cells (PBMCs) were isolated via density-gradient separation with Ficoll. Total CD8^+^ T cells were enriched by negative magnetic separation using the Miltenyi CD8^+^ T Cell Isolation Kit Human. Human polymorphonuclear leukocytes (PMNs) enriched pellets were diluted with PBS and 3% dextran solution in HBSS was added in a 1:1 ratio. After centrifugation, the supernatant was processed for red blood cell lysis according to manufacturer’s instructions (Red Blood Cell Lysis Solution, Miltenyi). Cells were either used fresh or cryopreserved according to standard procedures. Mouse neutrophils and CD8^+^ T cells were purified according to manufacturer’s instructions using anti-Ly6G and CD8^+^ T Cell Isolation Kit Mouse, respectively (Miltenyi Biotech). Purity of isolated human and mouse immune populations were checked by FACS (>95%).

### *In vitro* CD8^+^ T Cells and Neutrophils co-culture

CD8^+^ T cells activated with plate-bound anti-human CD3 (2mg/mL) and anti-human CD28 (2mg/mL) and freshly isolated neutrophils were co-cultured at different ratios in complete RPMI-1640 media.

### *In vitro* GZMK and SDF1 treatments

HT-29 and CACO2 cells were cultured in transwells in a ratio 1:9 (CACO2:HT-29) for 16 days when trans-epithelial electrical resistance (TEER) was measured to assess the epithelial formation. Cells were treated with rHu-GZMK 10uM for 24h. For treatment of neutrophils, freshly isolated cells were treated with rHu-GZMK 10uM or SDF1 100ng/ml and analyzed at different time points as indicated in the relative figure legend.

### Neutrophils migration assay

Neutrophil migration assays were performed using a custom-made microfluidic device in PDMS composed by three parallel channels (height 200µm, length 1.1cm, width central channel 1300µm, lateral channels 700µm) separated by two lines of trapezoidal pillars. The central channel was used to confine a collagen matrix gel, the right channel for cell injection and the left channel for treatments. PBS 10X, HEPES 80mM, NaOH 1N, 8.04mg/ml rat tail Collagen type I (Corning) and RPMI culture medium (without phenol red) were mixed on ice to obtain a collagen matrix solution. The collagen solution was injected in the central channel of the microfluidic chip and incubated at 37°C to allow the complete reticulation of the collagen matrix. The collagen matrix was hydrated by injecting culture medium into the lateral channels of the device. Culture medium was added to the reservoirs to avoid evaporation and chips were incubated at 37°C. Purified human neutrophils (3×10^6^/ml) from healthy donor were labeled with calcein-AM (green), Hoechst 33342 (blue), and DRAQ7 (red) and injected in the right channel of the microfluidic devices. The microfluidic chips were incubated at 37°C for 20 minutes on an angled surface (30 degrees) to allow neutrophils to lie at the interface between the lateral channel and the collagen gel. Chemoattractant solutions or culture medium were added to the left channel and time lapse experiments were performed using Nikon Eclipse Ti (3 fields for chip, 1 image every 2.30min for 4h at 10X magnification) to evaluate invasion of neutrophils in the collagen gel. Interstitial fluid from T and NAT of LN and HN patients, purified CXCL6 (400ng/ml), SDF-1 (100ng/ml) and IL-8 (100ng/ml) were tested. Migrating cells were counted and analyzed using ImageJ version 2.0.

### Neutrophils Gelatinase assay

The gelatinase activity was measured from control or SDF1 (100 ng/mL) treated neutrophils in triplicate, using DQ gelatin (molecular Probes, 100 µg/mL final concentration) following manufacturer’s recommendations. The fluorescent output was measured every 15 min at room temperature on a GloMax Discover (Promega Instrument) microplate reader (Ex= ∼495 nm, Em= ∼515 nm) and corrected for background. *Clostridium histolyticum* collagenase (0.15 U/mL) was used to generate an internal calibration curve.

### Neutrophils Adhesion assay

Purified human neutrophils were labeled with calcein-AM (green), Hoechst 33342 (blue) for 20 minutes at 37°C, as previously described (Htwe et al., 2018). After washing, neutrophils were incubated in RPMI media with or without SDF1 (100 ng/mL) for 1 hour at 37°C in a 96-well on a monolayer of human micro-endothelial cells (HMEC1). After two gentle washes with PBS to remove non-adherent cells, images of three fields were recorded for each well using an EVOS microscope (EVOS FL Cell Imaging System) at 4x magnification. Adherent neutrophils were measured by counting calcein positive neutrophils using ImageJ version 2.0.0.

### Multiparametric flow cytometry and sorting

High-dimensional flow cytometry was performed on T, NAT and PB. Conjugated antibodies used for flow cytometry are shown in Table 2. Briefly, single-cell suspensions were stained fresh or after thawing in pre-warmed RPMI with 10% FBS. Cells were washed in staining buffer (PBS with 2%FBS and 2mM EDTA) and incubated for 30min at 4°C with surface antibody cocktail in staining buffer, washed twice in staining buffer, fixed and permeabilized using Fixation/Permeabilization Solution Kit (BD, Cat # 554714), and stained intracellularly for 1h at 4°C with an antibody cocktail in the permeabilization solution. Samples were acquired using a FACSymphony A5 or a FACSCelesta BVYG equipped with FACSDiva software version 8.0.1 (all from BD Biosciences). Compensation Beads (ThermoFisher, Cat# 01-2222-41) were used to prepare single-stained controls for electronic compensation. Dead cells were excluded using Fixable Viability Stain (BD). To evaluate ex vivo cytokines production, CD8+ T cells were stimulated 3 h with PMA (20ng/mL), ionomycin (1ug/mL) and GolgiPlug protein-transport inhibitor (brefeldin A, 1:1000). CD8 T cell subsets were sorted to high purity using FACSAria III (BD Biosciences).

**Table 2.** Markers Multiparametric Flow Cytometry panel. Markers with the corresponding Fluorophore, Manufacturer and Catalog ID (Cat. #) are shown.

| Marker | Fluorophore | Manufacturer | Cat. # |
| --- | --- | --- | --- |
| Zombie Aqua |  | BioLegend | 423101 |
| TCRgd | PerCP-Cy5.5 | BioLegend | 331224 |
| NKG2A | FITC | Miltenyi | 130-113-568 |
| CD39 | APC-H7 | BioLegend | 328226 |
| TIGIT | APC | BioLegend | 372706 |
| CD25 | BV786 | BD | 741035 |
| CCR7 | BV711 | BD | 566602 |
| OX40 | BV650 | BD | 563658 |
| CD161 | BV605 | Biolegend | 339916 |
| CD27 | BV570 | BioLegend | 302825 |
| CD11b | BV510 | Biolegend | 301334 |
| PD1 | BV480 | BD | 566112 |
| CD103 | BV421 | BioLegend | 350214 |
| CD8 | BUV805 | BD | 564912 |
| CD28 | BUV737 | BD | 564438 |
| HLADR | BUV661 | BD | 565073 |
| CD4 | BUV615 | BD | 624297 |
| CD45RA | BUV563 | BD | 565702 |
| CD3 | BUV496 | BD | 564809 |
| CD69 | BUV395 | BD | 564364 |
| CD45 | PE-Cy7 | Biolegend | 304016304016 |
| CD56 | PE-CY5.5 | eBioscience | 35-0567-42 |
| CD127 | PE-CY5 | eBioscience | 15-1278-42 |
| CX3CR1 | PECF594 | Biolegend | 341624 |
| GZMB | APC-R700 | BD | 560213 |
| GZMK | PE | Santa Cruz | sc-56125 |

All the FCS files generated were analyzed and visualized with software FlowJo v10 (TreeStar). RPhenograph (Levine et al., 2015) package was used to analyze the multiparametric data from CRC patients derived T, NAT and PB. CD8 analysis was performed on 3000 events per sample after manual gating isolation of singlet, LD negative, CD45+CD3+CD8+ T cells. All samples were concatenated by the “cytof_exprsMerge function”. The number of nearest neighbors identified in the first iteration of the algorithm, K value, was set to 100. UMAP and tSNE representation were generated and visualized using FlowJo version 10 (FlowJo). Under-represented clusters (<0.5%) were discarded in subsequent analysis. The balloon plots, performed using the ggplots2 R package, generat a further metaclustering classification, interpolating MFI and frequency of each marker per cluster. MFI were multiplied to derive the integrated MFI (iMFI, rescaled to values from 0 to 100). Euclidean distance and Ward-linkage methods via cytofkit package were applied to generate hierarchical metaclustering of Phenograph generated clusters.

### Bead-based multiplexed ELISA

Multiplexed ELISA on interstitial fluid from matched T and NAT fresh tissues were performed on a Luminex 200 platform (Luminex Inc.,) using custom kits of pre-mixed antibody-coated beads (R&D System Inc., MN), which included the following analytes: CCL11_eotaxin, CCL13_MCP4, CCL17_TARC, CCL2_MCP1, CCL22_MDC, CCL26_EOTAXIN3,, CCL5_RANTES, CCL8_MCP2, CD25_IL2Ra, CX3CL1_FRACTALKINE, CXCL1_GROa, CXCL10_IP10, CXCL11_ITAC1, CXCL13_BLC_BCA1, CXCL4_PF4, CXCL5_ENA78, CXCL6_GCP2, EGF, IFNg, GMCSF, HGF, IL10, IL1ra_IL1F3, IL7, IL8_CXCL8, TRAIL, VEGFA, IL1b_IL1F2, IL5, IL6, IL-17F, IL22, IL23, TNFa, LXSAHM-01, CCL3_MIP1a, CCL4_MIP1b, FGFbasic_FGF2, GCSF, IL1a_IL1F1, IL2, IL4, IL-12 p70, IL13, IL15, IL-17/IL17A, CXCL12_SDF1, TGFB1, TGFB3. The assay was runned based on manufacturer reccomendations. Briefly, 50ul of samples together with kit standards were added to each well in duplicate and incubated with the diluted Microparticle Cocktail at 4C, ON, on a shaker at 850rpm. Unbound soluble molecules were removed by washing the plate. The Biotin-Antibody Cocktail specific to the analytes of interest was added to each well for 1h at RT. After washing again, the Streptavidin-Phycoeriythrin conjugated was added for 30 minutes at RT. After the final washing steps, the microparticles are resuspended in kit buffer and read on a Luminex 200 platform. The outputs (pg/mL) were visualized and statistically analyzed in R upon centering and scaling using the scale function in R (SD from mean pg/mL). Data were visualized together with sample annotations using the ComplexHeatmap package. Group comparisons were visualized as boxplots using the ggpubr package and statistically analyzed by applying the non-parametric Wilcoxon signed-rank test on the values, where significance is set at p-value < 0.05. Soluble molecule data were associated with other experimental data by Spearman correlation and visualized using the corrplot package.

### Immunohistochemistry (IHC) and Immunofluorescence (IF)

Slides (5um) from formalin-fixed paraffin embedded (FFPE) samples were processed for deparaffinization and rehydration. Heat-induced antigen retrieval (citrate buffer pH 6 or 9 – Thermofisher) was performed using the microwave. 3% H_2_O_2_ incubation was used for IHC preparation. Slides were incubated ON at 4°C with mouse anti-human CD66b antibody (BioLegend, 305102, 1:100 for both IHC and IF), anti-human CD8 (Invitrogen, 53-0008-82, 1:100), rabbit anti-human GRZMK (Invitrogen, LS-C119554-50, 1:100), rabbit anti-human E-cadherin (Abcam, Ab1416, 1:100), rat anti-human Ki67 (Ebioscience, SolA15, 1:200), rabbit anti-human EPCAM (Abcam, Ab32394, 1:400), mouse anti-human SDF1 (R&D, MAB350, 1:100), anti-human αSMA (Abcam, Ab8211, 1:200) in a blocking solution composed of 3% BSA, 5% goat serum (Invitrogen, 10000C) and 0,1% Triton in PBS. After incubation with HRP-secondary antibody (30min) and Aminoethyl Carbazole (AEC) + High Sensitivity Substrate Chromogen (Dako) (for IHC) or fluorophore conjugated antibodies (Invitrogen, A32723, goat anti-mouse 488; Invitrogen, A21247, goat anti-rabbit 647; Invitrogen, A21206, goat anti-rabbit 488; Invitrogen, A21424, goat anti-mouse 555; 1:500; Invitrogen, A21434, goat anti-rat 555, all 1:500 for 1 hour at RT), slides were mounted and acquired by using Aperio or SP8 confocal microscope (Leica), respectively.

### Imaging analysis

For quantification of GZMK signal on human colorectal cancers, sections were labeled with DAPI, anti-GZMK, anti-CD66b and Alexa Fluor 488-conjugated anti-CD8 primary antibodies and images captured with a Nikon CSU-W1 spinning disk (Nikon Europe BV) using a 40X/1.15 NA water immersion objective lens (pixel size 0.1625 x 0.1625 um^2^). For the tumor area identification, previews of tissue sections on the DAPI channel were acquired with a 10X/0.3 NA dry objective lens. Inside the tumor areas, 3 to 4 regions were randomly chosen for the acquisition with a 40X/1.15 NA water immersion objective lens. Each region was made of 12 adjacent tiles covering a total area > 4 mm^2^ per tissue section.

The quantification of the GZMK signal in CD8^+^ cells was done with a custom-made Fiji macro (Schindelin et al., 2012). Briefly, images were pre-processed with denoising and background subtraction, and nuclei were segmented on the DAPI channel with the plugin StarDist (Schmidt et al., 2018), using the built-in model Versatile (fluorescent nuclei) and the intensity parameters, including “mean intensity” and “raw integrated density”, of CD8, GZMK and CD66b fluorescence were quantified in a band of thickness 1 um around each nucleus.

Statistical analysis was performed in R. CD8^+^ cells were identified from the database by setting criteria on the intensity parameters of the specific fluorescence. Particularly, a cell was considered as positive for CD8 fluorescence channel if its “raw integrated density” was above a threshold value, set by the comparison in images of the “raw integrated density” values of cells expressing CD8 fluorescence and the “raw integrated density” of cells expressing only autofluorescence signal. Moreover, a further correction regarding the cells that had the band around the nucleus overlapping with red blood cells was applied: the ratio of the “raw integrated density” in the green channel versus the red channel was computed and only cells with a ratio green/red > 4 were considered as positive for CD8. The same procedure described above was used for the detection of CD66b positive cells (for red blood cells correction a ratio red/green > 2 was set to consider cells as CD66b positive).

Differences between two groups were calculated by unpaired, two-tailed Student’s t test. The analysis of GZMK fluorescence in CD8+ cells lying in the proximity of a neutrophil (CD66b^+^) was performed as follows: the distance between a CD8^+^ cell and its nearest neutrophil was computed per each field of view. Only distances below 10 um were taken into account to select pairs of CD8+ cell-neutrophil that were close to each other. The mean intensity of GZMK fluorescence in CD8+ cells close to neutrophils was compared to the GZMK mean intensity of cells close to neutrophils that were not positive for CD8.

For the quantification of SDF-1 in cells positive for αSMA, human colorectal cancers sections labeled with DAPI, anti-SDF-1, anti-αSMA and anti-EPCAM primary antibodies, and captured with a Nikon Eclipse Ti widefield microscope (Nikon Europe BV) using a 20X/0.75 NA dry objective lens (pixel size 0.32 x 0.32 um^2^). For the tumor area’s identification, previews of tissue sections were acquired with a 4X/0.2 NA dry objective lens. Inside the tumor areas, regions were randomly chosen for the acquisition with 20X/0.75 NA dry objective lens for a total acquired area > 35 mm^2^ per tissue section.

The quantification of SDF-1 signal in αSMA^+^ cells was done with Qupath v. 0.2.3 (Bankhead et al., 2017) through a custom-made script. Briefly, cell nuclei were detected on DAPI channel with the StarDist extension and a cell expansion of 1 um was applied in order to measure fluorescence in the cytoplasmic/membrane cell’s area. For the nuclei segmentation with StarDist, a custom model was trained and tested using the StarDist (2D) network in ZeroCostDL4Mic (von Chamier et al., 2021): a training dataset was generated by the manual annotation of nuclei in QuPath and labeled images were exported for the deep-learning training; the model’s training and quality control assessment were performed with the StarDist (2D) notebook. For the detection of pixels positive for αSMA staining, a pixel classifier was created with the QuPath’s function Create thresholder and the quantification of the positive area inside each segmented cell was added as a measurement parameter in the detection’s results table. Other two-pixel classifiers in the red and far-red channels were similarly created to be used in the statistical analysis step. The expression of SDF1 was quantified as the mean intensity of the corresponding fluorescence signal in the band around the nucleus.

Statistical analysis was performed in R. Cells with an area > 3 um2 (estimated as the 10% of the area of a band created around a nucleus with area size 46 um2) of pixels classified as positive for αSMA were considered as αSMA+. Cells overlapping with red blood cells were discarded from the analysis when presenting positive area > 0 um2 for the pixel classifiers in green, red and far-red channels. Differences between two groups were calculated by unpaired, two-tailed Student’s t test.

### scRNA-seq

T and NAT single cell suspension after counting was resuspended in PBS without Ca2+ and Mg2+ with 0.04% BSA. Approximately 2,000 cells/ul from each sample were used for the analysis. Briefly, every sample was loaded into one channel of Single Cell Chip A using a Chromium Single Cell 3′ v2 Reagent Kit (10x Genomics). After capture and lysis, complementary DNA was synthesized and amplified over 14 cycles according to the manufacturer’s protocol (10x Genomics). Libraries were prepared from 50 ng amplified cDNA. Sequencing was performed using a NovaSeq 6000 System (Illumina). An average sequencing depth of at least 50,000 reads per cell was obtained for each sample.

### scRNAseq analysis

Single cells of samples from 11 patients (each one sequenced in the far tumor (D) and tumor site (T)) were sequenced with the 3’ v3 chemistry and aligned using Cellranger count v. 3.1 on human genome hg38. Only droplets with a minimum number of unique molecular identifiers (UMI) were considered as “cells”; this was done by default by the Cellranger program.

We carried out a preliminary analysis using data of all the samples collected from the 11 patients (D and T, total of 89715 cells, data not shown). Based on total cells in each patient and the distribution of each patient’s cells within groups identified by UMAP, we excluded patients 3, 8 and 6 from further analyses. Thus, 52316 cells were processed.

We used the Seurat v. 3 R package to analyse single cell data (Stuart et al., 2019). To identify cell populations or sub-populations, uniform manifold approximation and projection (UMAP) was carried out from the first principal components of the PCA based on the expression of the most variable genes, normalized using SCT transformation as suggested in Seurat documentation (“SCTransform” function, (Hafemeister and Satija, 2019). The different clusters in UMAPs were found using nearest neighbor clustering (SNN) with the function “FindClusters” of the Seurat package. In particular:

a. For global cell population (52316 cells, fig. Supp. 6A): for UMAP, the first 30 principal components were used, and n.neighbors=60, min.dist=0.005 were set. For SNN clustering, top 20 principal components were used, with other parameters k.param=10, prune.SNN=1/10. The resolution used in the “FindClusters” function was set to 0.11.
b. For T/NK cells (8716 cells, fig. Supp. 6E): cells belonging to the “T/NK’’ cluster of the previous step (a) were used. The UMAP was built using the first 30 principal components and n.neighbors=40, min.dist=0.002 were set as parameters. SNN clustering was carried out as above (a) but with a resolution of 0.09 in the “FindClusters’’ function. Finally, cells with the first UMAP dimension (UMAP_1) > 8 were considered as outliers and removed.
c. To further characterize T CD8+ cell subtypes (fig. 6A), cells belonging to the clusters 3 and 4 were extracted from the previous step (b) and the UMAP was built using the first 15 principal components from the expression of the following cell-specific markers that presumably would identify the expected subtypes: TRDC, PRF1, CD3D, CD3E, TRGC1, TRGC2, LEF1, IL6R, NOSIP, SELL, CCR7, CD4, BATF, TIGIT, FOXP3, TNFRSF4, CD8A, CD8B, GZMH, FGFBP2, GZMB, GNLY, ZEB2, HLA-DRB5, CX3CR1, GZMA, IFNG, CCL4, SBF2, KLRB1, LTB, IL7R, FOS, CD40LG, DPP4, IRF1, GZMM, GZMK, HLA-DRB1, HLA-DRA, CCL5, CTLA4, PDCD1, HAVCR2, LAG3, CD38, ENTPD1, LAYN, CXCL13, TOX. Other parameters for the UMAP were n.neighbors=20, min.dist=0.01. For SNN clustering, the top 15 principal components were used, with other parameters being k.param=10, prune.SNN=1/10. The resolution parameter in the “FindClusters” function was set to 0.5.

Differential expression of genes between different cell subgroups was carried out with the “FindMarkers” function in the Seurat package, using the Wilcoxon test. Thresholds were set to log2FC=0.25 and adjusted p value=0.05. For Figures 6C, 6F and Supp. 7D and E the size of the points corresponds to the score of the specific signature; this score is defined, for each cell, as the sum of the counts of the genes belonging to a specific signature / the total sum of the counts of that cell. The points with the biggest size correspond to a score higher than 80% of the distribution of the scores of all the cells.

Cell types in the entire population (Fig. Supp 6A) were identified with the help of scMCA (single cell mouse cell atlas) package (Sun et al., 2019) and then evaluating the expression of the cell type specific markers. Pseudotime analysis (fig. 6E) was done using Monocle v. 3.

### *In vivo* animal experiments

C57BL/6J were subcutaneously injected with 1 × 10^6^ MC38 cells/mouse. Animals were euthanized when the tumor reached the volume of <1cm^3^ and >1cm^3^ to evaluate CD8^+^ T cell and neutrophil infiltration during progression. Blood was collected and spleen and tumors harvested and dissociated to a single cell suspension. Tumors were digested (1,5 mg/mL collagenase, 0,75 mg/mL hyaluronidase and 0,1 mg/mL DNaseI) in RPMI 10% FBS at 37°C. Cell suspension was filtered through 70-μm cell strainers, washed, counted and stained for multiparametric flow cytometry.

### Survival analysis methods

We investigated the interplay between immune cytolytic activity and neutrophils infiltration on overall (OS) and disease-free survival (DFS) in the colon adenocarcinoma cohort (COAD) of The Cancer Genome Atlas (TCGA) (Weinstein et al., 2013) by performing a Kaplan-Meier analysis. The cytolytic activity index was quantified by computing the geometric mean of granzyme A and perforin expression (Rooney et al., 2015) and samples were split according to the Maximally Selected Ranks Statistics cutoff (“maxstat”, version 0.7-25). Neutrophils relative abundance was obtained by running CIBERSORTx (Rooney et al., 2015) with a validated leukocyte gene signature matrix (LM22) on RNAseq data of COAD samples. Samples were divided into two groups according to the presence or absence of Neutrophils. Similarly, CIBERSORTx-derived gene expression signature was employed to infer the GZMK^high^ cell subtype’ abundance and analyze its association with survival. First, the single-cell raw counts matrix of CD8^+^Tem population was split into two subgroups expressing “high” or “low” values of GZMK and used <u>it</u> to fe<u>e</u>d CIBERSORTx, building a gene signature able to discriminate the two cell sub-populations. Secondly, we collected survival tables of TCGA-COAD and TCGA-LUAD cohorts from cBioPortal (Cerami et al., 2012) and ran Kaplan-Meier analysis stratifying patients with “high” and “low” infiltration of CD8^+^ Tem cells expressing high levels of GZMK. Survival analyses were carried out on R software version 4.0.5), by using “survival” (version 3-2-11) and “survminer” (version 0.4.9) packages.

### Statistics and Data visualization

Results were analyzed and visualized on Prism version 9.2.0 (GraphPad). All statistical analyses were conducted using GraphPad Prism version 9.2.0 or R software version 4.0.2. Differences between two groups were calculated by two-tailed Student’s t test. Multiple comparisons were performed by one-way ANOVA followed by Bonferroni’s multiple comparison tests. Significance was set at P values ≤ 0.05. For all figures: *, P ≤ 0.05–0.01; **, P ≤ 0.01–0.001; ***, P ≤ 0.001–0.0001; ****, P ≤ 0.0001. Unless noted in the figure legend, all data are shown as mean ± SEM. The n numbers for each experiment and the numbers of experiments are noted in the figure legends.

## RESULTS

### Early-stage CRC patients can be stratified based on the abundance of CD15^high^ neutrophils in the tumor microenvironment

Despite consistent T cell infiltration, over a third of early-stage CRC patient relapse, suggesting that the anti-tumor immune response is somehow “leaky”. In this regard, characterizing the heterogeneity of immune cells within and across CRC patients would be essential to fully understand the mechanisms behind immune evasion. We profiled peripheral blood (PB), tumor (T) and normal adjacent tissue (NAT >10 cm from tumor tissue) from treatment-naïve patients (n= 42) with early-stage CRC (8^th^ ed. AJCC stages I, II and III) undergoing surgical resection (Figure 1A). Details of all patients involved in the study are summarized in Table 1. Among the infiltrating innate immune cell populations, we observed no significant difference on dendritic cells (DCs), lower proportion of NK cells in T compared to NAT and enrichment in macrophages in T, albeit they were present at low frequencies (Figure Supp. 1A). On the other hand, neutrophils were higher in T compared to NAT both in number and frequency, as assessed by IHC (CD66b^+^, Figure Supp. 1B) and FACS (Figures 1B) (Bronte et al., 2016; Gustafson et al., 2015), respectively. Neutrophils were identified as CD45^+^CD11b^+^CD15^+^ cells along with intermediate/low expression of CD33 and negative staining for HLA-DR (Figure Supp. 1C) (Gustafson et al., 2015). Analysis of CD15 expression on the surface of intratumor neutrophils (Pillay et al., 2013) revealed two distinct neutrophil populations identified by high and low levels of CD15 (referred to as CD15^high^ and CD15^low^, respectively, Figure Supp. 1D). Despite being overall similar in NAT, CD15^high^ neutrophils were enriched in tumors in a fraction of patients (Figure Supp. 1E). Thus, based on the CD15^high^ and CD15^low^ neutrophil populations, we divided our cohort in 2 groups, namely low (LN) and high CD15^high^ (HN) neutrophils (Figures 1C and D).

**Figure 1.**
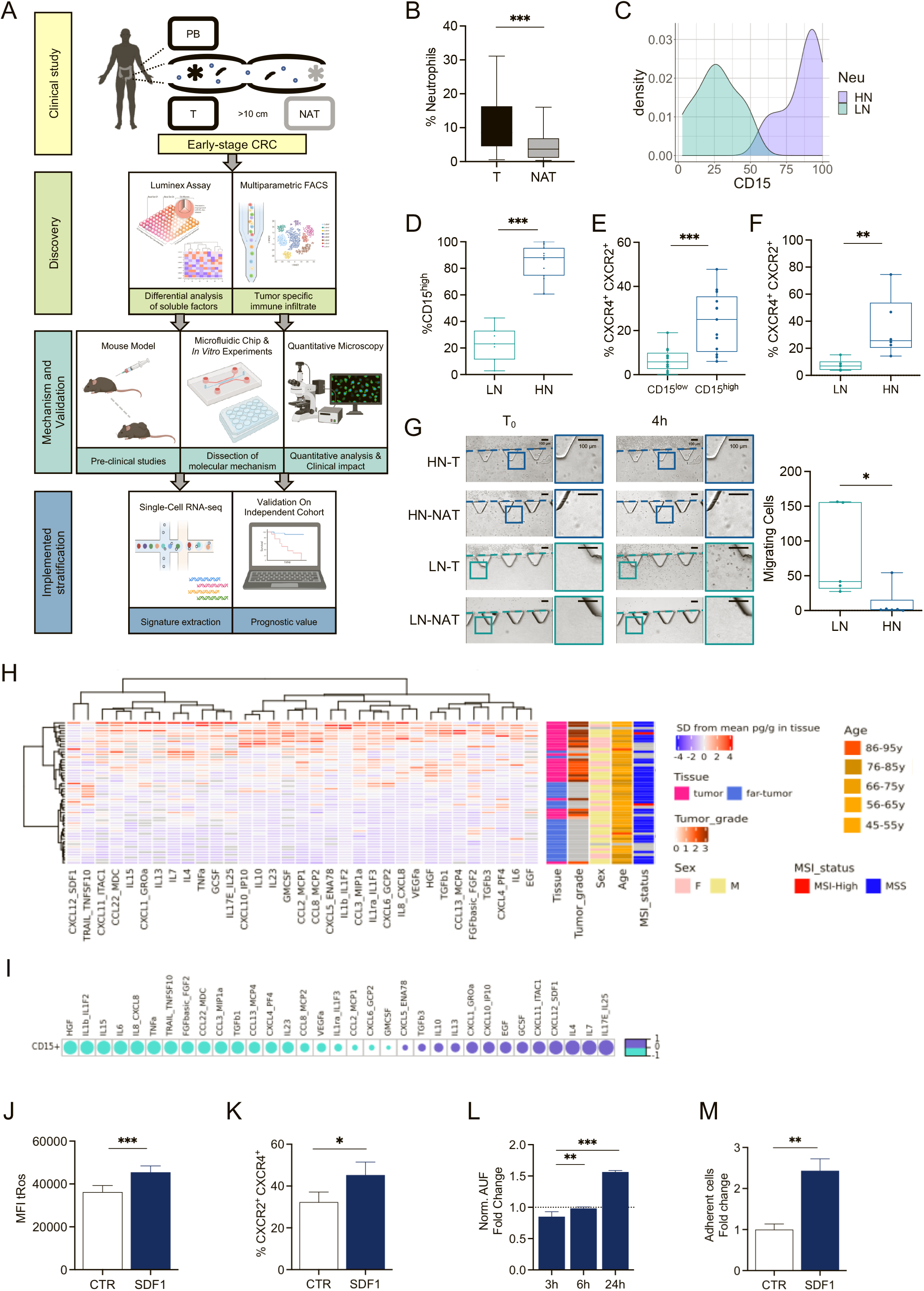
Early-stage colorectal cancer (CRC) patients can be stratified based on the abundance of CD15^high^ neutrophils in the tumor microenvironment (TME). **A.** Study design. Immune analysis of Tumor tissues (T), Normal Adjacent Tissues (NAT) (10 cm from T) and Peripheral Blood (PB) of early-stage CRC patients (n=42). **B.** Frequency of Tumor Associated Neutrophils (TAN) defined as CD45^+^ CD11b^+^ HLADR^−^ CD56^−^ CD66b^+^ within CD45^+^ immune compartment in T compared with NAT samples in CRC patients (total n=22). Bars indicate 90-10 percentile. **C.** Density plot of CD15 expression within TAN defining a bimodal distribution in the CRC cohort of patients. Patients were clustered as low (LN) or high (HN) based on CD15 expression. **D.** Frequency of CD15^high^ neutrophils in LN and HN patients. Bars indicate 90-10 percentile. **E.** Bar plot of CXCR4^+^ CXCR2^+^ cell frequency within CD15^low^ and CD15^high^ neutrophils. **F.** Bar plot of CXCR4^+^ CXCR2^+^ cell frequency within LN and HN neutrophils. **G.** Representative brightfield images of neutrophils migration assay on the microfluidic device. Interstitial Fluid of T and NAT from HN and LN patients were tested. Bar plot of number of migrating neutrophils on chip upon stimulation with IF from HN (N=6) and LN (N=5) patients. Scale bar 100mm. **H.** Heatmap showing the abundance of soluble molecules in from IF of T (tumor, light blue) and NAT (far-tumor, pink) tissues. For all the samples, tissue type, tumor grade, sex, age and Microsatellite Instability (MSI) status are indicated. **I.** Spearman correlation between soluble molecules in T IF and CD15^high^ neutrophils. **J.** Frequency of CXCR4^+^ CXCR2^+^ in Healthy Donors (HD) fresh-isolated neutrophils treated with SDF-1. **K.** Bar plot representation of total ROS (tRos) MFI within SDF-1 treated-neutrophils from healthy donors (HD). **L.** Gelatinase activity assay on neutrophils freshly-isolated from HD and treated with SDF-1 for the indicated time. Bar plot representation of fold change over untreated control. **M.** Quantification of adherent neutrophils on endothelial cells after 2 hours treatment with SDF-1.

In line with previous studies, our data suggested that CD15^high^ neutrophils, which predominantly infiltrate HN tumors, were in a N2-like tumor-promoting functional state (Fridlender et al., 2009). Indeed, we reported that CXCR2^+^CXCR4^+^ double positive cells were significantly higher in the CD15^high^ compared to the CD15^low^ neutrophil compartment (Figure 1E). Accordingly, HN tumors showed increased frequencies of N2-like tumor-promoting CXCR2^+^CXCR4^+^ neutrophils (Figure 1F). In line with a tumor-promoting phenotype, CD15^high^ neutrophils expressed significantly higher levels of CD10, a cell surface marker that was proposed to distinguish T-cell suppressive from T-cell stimulatory neutrophils (Figure Supp. 1F) (Fridlender et al., 2009; Marini et al., 2017). Moreover, exposure of neutrophils to interstitial fluid from HN tumors induced higher production of total reactive oxygen (tROS) species (Figure Supp. 1I). Of note, the frequencies of CXCR4^+^CXCR2^−^ neutrophils, previously defined as aged (Jaillon et al., 2020a), were comparable between CD15^high^ and CD15^low^ neutrophils as well as in HN and LN tumors (Figures Supp. 1G-H). Overall, these data revealed that CRC patients can be stratified based on the abundance of a CD15^high^ neutrophil population, which might have tumor promoting functions.

Neutrophil homeostasis in peripheral tissues results from the balance between clearance by macrophages and recirculation regulated by soluble factors (Cox et al., 1995; Grigg et al., 1991) (Buckley et al., 2006; Robertson et al., 2014; Wang et al., 2017; Woodfin et al., 2011). Indeed, frequency of macrophages anticorrelated with CD15^high^ neutrophils (Figure Supp. 1J) and was significantly lower in HN compared to LN (Figure Supp. 1K), suggesting that lower clearance in HN tumors might contribute to higher abundance of neutrophils, as previously reported (Peng et al., 2021a).

Next, we asked which was the contribution of soluble chemoattractants in determining the CD15^high^ neutrophil difference between HN and LN. This was tested on a microfluidic device that enabled measurement of the ability of freshly isolated neutrophils to migrate through a 3D collagen matrix. Surprisingly, migration was higher when neutrophils were exposed to LN compared to HN tumor interstitial fluids (Figures 1G), suggesting that the higher CD15^high^ neutrophils observed in HN compared to LN did not reflect an increase in their recruitment into the TME. On the contrary, exposure of neutrophils to interstitial fluid from HN tumors increased expression of CD15 when compared to LN (Figures Supp. 1L), suggesting an active role of the TME in driving neutrophil function. Therefore, we profiled 49 inflammatory soluble factors in the interstitial fluid of T and NAT from CRC patients, of which 33 gave consistently measurable results (Figure 1H). While not significantly associated neither with demographics nor tumor grade, we found out that a subset of these factors was directly correlated with the frequency of CD15^high^ neutrophils (Figure 1I). In particular, CD15^high^ neutrophils were positively correlated with IL-17E/IL-25, which plays important roles in inflammation occurring in the gut (Caruso et al., 2009; Owyang et al., 2006; Wang et al., 2014), it has been shown to upregulate the expression of CXCL1 and CXCL10 (Senra et al., 2019) and promotes a Th2-like responses by inducing IL-4, IL-5 and IL-13 expression (Fort et al., 2001). In agreement, we found that CXCL1, CXCL10, IL-4 and IL-13 were positively correlated with CD15^high^ neutrophils. Interestingly, IL-4/IL-13–stimulated neutrophils have been reported to have diminished ability to form NETs and migrate toward CXCL8 (Impellizzieri et al., 2019), suggesting that neutrophils from HN and LN might be functionally different. Enrichment of G-CSF and IL-10 in HN further supported the role of the TME in driving neutrophil polarization toward N2-like functional state (Ferrante; Fridlender et al., 2009; Gerlini et al., 2004; Gholamin et al., 2009; Joshita et al., 2009; Kyo et al., 2000; Moore et al., 2001). Among the detected factors, CXCL12/SDF-1 (from now on referred to as SDF-1) was of particular interest because of its role in neutrophil trafficking and retention at inflammatory site through its binding to CXCR4 (C et al., 2003; Filippo and Rankin, 2018), which we found to be highly expressed on CD15^high^ neutrophils together with CXCR2 (Figures 1E-F). Imaging analysis confirmed that overall levels of SDF-1 were higher in HN compared to LN tumors and this was due to the combination of a larger number of αSMA^+^ cancer associated fibroblasts in HN and their higher expression of SDF-1 (Figure Supp.1M-N).

Despite surface expression of CXCR4, SDF-1 treatment did not foster neutrophil’s migration *in vitro.* On the contrary, IL-8 and CXCL6, well-known chemo-attractants found to negatively correlate with CD15^high^ TANs *in vivo*, showed a pro-migratory effect (Figures Supp. 1O-P). On the other hand, neutrophils treated *in vitro* with SDF-1 showed higher frequency CXCR2^+^CXCR4^+^ double positive cells (Figures 1J and Supp. 1Q-R), an elevated level of total ROS (Figure 1K), produced more gelatinase (Figure 1L) and adhered more to micro-endothelial cells compared to untreated controls (Figure 1M), which recapitulates, at least in part, *in vivo* observations. The slight increase in CD62L upon SDF-1 treatment supported the idea that these neutrophils were not aged (Figure Supp.1S) (Peng et al., 2021b). In summary, these results showed that tumor infiltration of CD15^high^ neutrophils distinguished between LN and HN patients, and suggested that elevated concentrations of SDF-1 changed neutrophil functional state rather than their recruitment.

### The TME of CRC is infiltrated by two main CD8^+^ T cell populations identified by differential expression of CD39 and GZMK

We next hypothesized that the different abundance of CD15^high^ neutrophils might alter the immune landscape in CRC TME and, in particular, CD8^+^ T cells. CD8^+^ T cell infiltration is unanimously accepted as a favorable prognostic marker in CRC (Chiba et al., 2004; Galon et al., 2014; Naito et al., 1998; Pagès et al., 2009b; Prall et al., 2004; Zlobec et al., 2010). However, querying the collection of non-metastatic CRC patients in the TCGA database with a cytolytic CD8^+^ T cell transcriptional signature did not result in an effective stratification (DFS, p=0.12 Figure Supp. 2A), suggesting that other factors might influence CD8^+^ T cell-mediated response and contribute to determining the clinical outcome. Therefore, we employed high dimensional flow cytometry to carry out a detailed assessment of the diversity in the CD8^+^ T cell compartment, including markers of memory and effector differentiation (CD45RA, CCR7, CD27, CD28, CD127, CX3CR1, CD161), activation (OX40, CD25, HLA-DR), inhibitory receptors (PD1 and TIGIT), tissue residency and tumor reactivity (CD69, CD103 and CD39) and effector molecules (GZMB and GZMK). The whole list is reported in Table 2. Within the TME, CD3^+^ T cells represented over 60% of the immune infiltrate (Figure Supp 2B) and were enriched in T compared to NAT across patients (Figure 2A). In the T, the composition of the CD3^+^ T cells included CD4^+^ Th (T_Th_) cells, CD8^+^ T cells, T regulatory (T_Reg_) cells and T gamma-delta (Tγδ) cells, with CD4^+^ Th and CD8^+^ T cells being predominant (Figure 2B).

**Figure 2.**
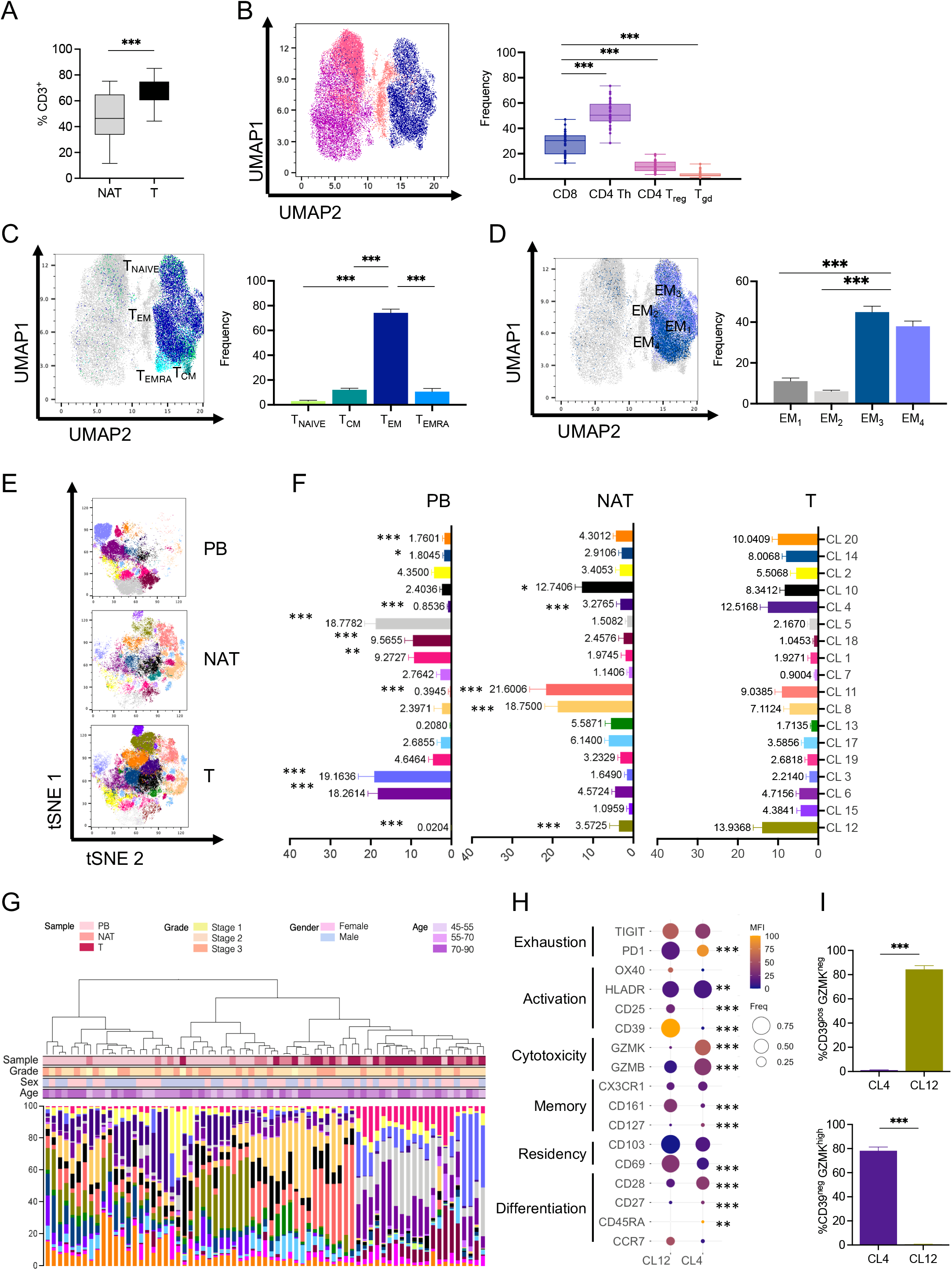
High-dimensional single cell analysis of CD8^+^ T cells identifies a tumor-specific CD39^−^ GzmK^+^ population enriched in early-stage CRC. **A.** CD3 Lymphocytes frequency within CD45^+^ in T compared with NAT. Bars indicate 90-10 percentile. **B-D.** UMAP Representation of concatenated CD3^+^ T cells from T sample with bar plot quantification. **B.** Cytotoxic CD8^+^ T cells (blue), Th CD4^+^ (purple), Treg CD4^+^, CD25^+^, CD127^−^ (pink) and γδT cells (orange) distribution. **C.** Naive T cells (light green), central memory T cells (dark green), effector memory T cells (dark blue), effector memory cells re-expressing CD45RA (light blue) distribution. **D.** effector memory T cells subtypes (EM_1-4_), see main text for description. **E.** tSNE representation of Phenograph algoritm identified 18 clusters of concatenated CD8+ T cells (3,000 cells/sample) from T (n=34), NAT (n=17) and PB (n = 22) samples from CRC patients. **F.** Cluster representation in different tissues: PB, NAT, T. **G.** Hierarchical metaclustering of CRC patients derived samples based on Phenograph identified clusters using Ward’s minimum variance method. For all the samples, sample type, grade, sex, age are indicated. **H.** Balloon plot of the average expression levels and expression frequencies of marker of differentiation, residency, memory, cytotoxicity, tumor reactivity, activation and exhaustion in clusters identified in (E-F) and enriched in T. **I.** Bar plot of CD39^+^ GzmK^−^ and CD39^−^ GzmK^+^ frequency within Cluster 4 and Cluster 12, respectively. *, P < 0.05**, P < 0.01 ***, P < 0.001.

Focusing on the CD8^+^ T cell compartment, naive (T_N_ identified by CD45RA^+^CCR7^+^) and central memory (T_CM_ identified by CD45RA^−^CCR7^+^) CD8^+^ T cells were rare both in the T and in NAT compared to PB (Figure Supp. 2C). In line with previous reports (Pagès et al., 2009b), the majority (about 80%) of the infiltrating CD8^+^ T cells were effector memory cells (T_EM_), characterized by CCR7^−^CD45RA^−^ expression (Figures 2C and Supp. 2C). Therefore, based on differential CD27 and CD28 expression (Figure Supp. 2D), we further divided the T_EM_ into 4 phenotypically and functionally distinct subpopulations called EM_1_ (CD27^+^CD28^+^), EM_2_ (CD27^+^CD28^−^), EM_3_ (CD27^−^CD28^−^) and EM_4_ (CD27^−^CD28^+^) (Romero et al., 2007). The T_EM3_ and T_EM4_ T cell compartment, which represented effector-like and memory-like phenotypes respectively, were the most represented in T (Figures 2D). We detected low frequencies of terminally differentiated effector memory cells re**-**expressing CD45RA (T_EMRA_) (Figures 2C and Supp. 2C), a phenotype associated with terminal T cell differentiation (Thome and Farber, 2015). In accordance, only a minority of CD8^+^ T cells exhibited traits that have been previously associated with “T cell exhaustion” (McLane et al., 2019), such as co-expression of PD-1, LAG3 and TIM3 (Figure Supp. 2E).

Analysis with t-distributed stochastic neighbor embedding (tSNE) confirmed that CD8^+^ T cell profiles were substantially different in the PB, NAT and T compartments also at single-cell level (Figures 2E-F), independently from other factors like patient’s age, gender or tumor stage (Figure 2G). Based on the calculated mean fluorescence intensity (MFI) and frequency of different markers (Figures Supp. 2F-G), we identified 20 different clusters (CL) in the CD8^+^ T cell compartment (Figures Supp. 2G-H, CL 9 and 16 were excluded from further analyses because their frequencies were below 0.5%). A number of clusters were enriched in PB compared to NAT and T and were characterized by lack of expression of residency and activation markers (CD69, CD103 and CD39). These included: CL3 - annotated as naïve (T_N_) and characterized by high levels of CD45RA, CCR7 and CD127 expression and lack of activation markers; CL1, 5 and 18 were annotated as T_EMRA_ being characterized by low CCR7 level, general lack of expression of memory markers along with high levels of the cytolytic molecule GRZB; CL6 - annotated as central memory (T_CM_) because of the expression of several memory-associated markers (CCR7, CD127, CD27, CD28) but absence of activation and cytotoxic markers (Figures 2H and Supp. 2G). Four other clusters, instead, were enriched in T and NAT compared to PB, indicating that they were tissue-infiltrating CD8^+^ T cell subsets. These were: CL14 and 20 mainly composed by CD45RA^neg^CCR7^neg^CD28^high^CD27^low^CX3CR1^pos^ T_EM_ cells, and CL8 and 11 annotated as tissue-resident memory T cells (T_RM_) since they displayed high expression of markers of memory (CD127 and CD28) and tissue residency (CD69 and CD103). These two tissue-infiltrating T_RM_ subsets expressed intermediate levels of CD161 and were mostly present in NAT (Figures 2F and Supp. 2G). In line with previous reports (Fergusson et al., 2015), this population appeared to be different from the CD161^high^ mucosal associated invariant T cells (MAIT) in CL19, which appeared to be equally represented in all three different compartments in early CRC patients.

We next focused on CL12 and CL4, which were specifically enriched in T compared to NAT and PB, and found that they were mainly composed by a T cell population reminiscent of CD8^+^ T_EM_ cells (CD45RA^low^CCR7^low^CD28^int^CD27^low^CD8^+^). They showed low or intermediate levels of CD127 and CX3CR1, and differential expression of activation, exhaustion and cytotoxicity markers (Figures 2H and Supp. 2G). In particular, CL12 identified CD8^+^ T cells with negligible expression of GZMK and high levels of CD39, a marker that has been recently associated with tumor antigen encounters (Duhen et al., 2018; Simoni et al., 2018). On the contrary, CL4 was primarily composed by CD8^+^ T cells characterized by lack of CD39 and elevated levels of GZMK (Figure 2I). Collectively, these data showed that CRC TME is mainly infiltrated by two distinct CD8^+^ T_EM_ cells populations that could be distinguished based on GZMK and CD39 expression (GrzK^high^CD39^neg^ and GrzK^low^CD39^pos^).

Using a manual gating strategy, we observed that CD8^+^ T_EM_ compartment was predominantly occupied by CD39^neg^ T cells (Figure 3A). However, the expression of CD69 and CD103 in the CD39^neg^ CD8^+^ T_EM_ cells indicated that they were not derived from blood contamination (Figures Supp. 3A-B). Thus, we decided to focus our attention on the characterization of this newly emerged population of CD8^+^ T_EM_, annotated as CL4 and identified by the unique expression of CD39 and GZMK. Manual gating analysis showed that GZMK^high^ CD8^+^ T_EM_ cells were almost entirely CD39^neg^ (Figure 3B) and were characterized by significantly lower expression of PD1 and higher GZMB when compared to their GZMK^low^ counterpart (Figure 3C). When FACS-sorted based on CD39 expression and assessed for their ability to produce effector molecules, CD39^neg^ CD8^+^ T cells produced significantly higher levels of GZMK and retained the ability to produce IFNγ, TNFα and GZMβ compared to their CD39^high^ counterparts (Figure Supp. 3D). In conclusion, we reported that the TME in early-CRC patients was highly infiltrated by a novel GZMK^high^ CD8^+^ T_EM_ population, which lacked CD39 expression and was distinct from the “exhausted-like” GZMK^+^ CD8^+^ T cell subpopulations recently reported by others (Hornburg et al., 2021; Mogilenko et al., 2021; Zhang et al., 2018).

**Figure 3.**
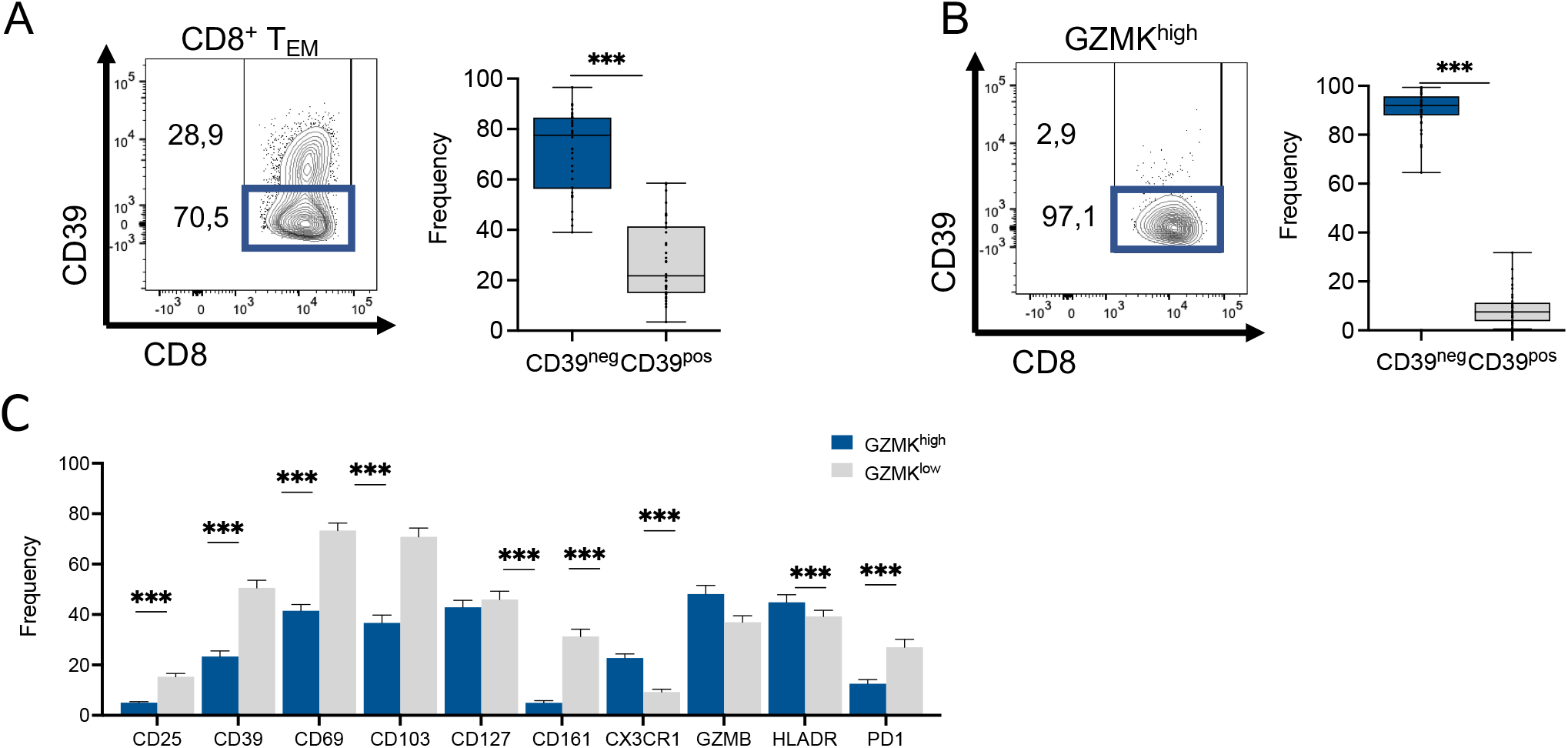
Phenotypic characterization of the CD39^−^ GZMK^+^ CD8^+^ T cell population infiltrating early stage CRC patients. **A.** Representative contour plot and quantification of CD39^−^ and CD39^+^ frequency within CD8^+^ T. **B.** Representative contour plot and quantification of CD39^−^ and CD39^+^ frequency within GZMK^+^ CD8^+^ T. **C.** Bar plot showing frequencies of the indicated markers within GZMK^+^ (dark blue) and GZMK^−^ (light gray) CD8^+^ T. ***; P < 0.001.

### Neutrophils modulate CRC infiltrating CD8^+^ T cells increasing their GZMK expression

Neutrophils influence CD8^+^ T cell responses both in animal models and humans. However, evidences are still sparse and often contrasting (Eruslanov et al., 2014; Jaillon et al., 2020a; Li et al., 2020; Sangaletti et al., 2014a), and even the stratification of non-metastatic CRC patients in the TCGA database was not effective when combining neutrophils to the cytolytic CD8^+^ T cell transcriptional signature (Figure Supp. 4A, p=0.44). Thus, since CRC patients can be distinguished in two distinct subgroups based on the abundance of CD15^high^ neutrophils, we sought to better understand their link with CD8^+^ T cells. Strikingly, Spearman correlation analysis revealed that frequencies of intra-tumor CD15^high^ neutrophils were positively correlated with frequencies of GZMK^high^ CD8^+^ T_EM_ cells (Figure 4A). Of note, no significant correlation was found between CD15^high^ neutrophils and other CD8^+^ T cell differentiation states or activation markers (Figures Supp. 4B and C). Since CD4^+^ T cells were the predominant population in the CD3^+^ T cell compartment, we sought to address their potential contribution to GZMK production. Our data showed that CD4^+^ T cells produced low amounts of GZMK and they were neither correlated with CD15^high^ neutrophils abundance nor differentially present in the TME of HN and LN (Figures Supp. 4D-F). Together, these data prompted us to investigate the specific cross-talk between CD15^high^ neutrophils and GZMK^high^ CD8^+^ T_EM_ cells.

**Figure 4.**
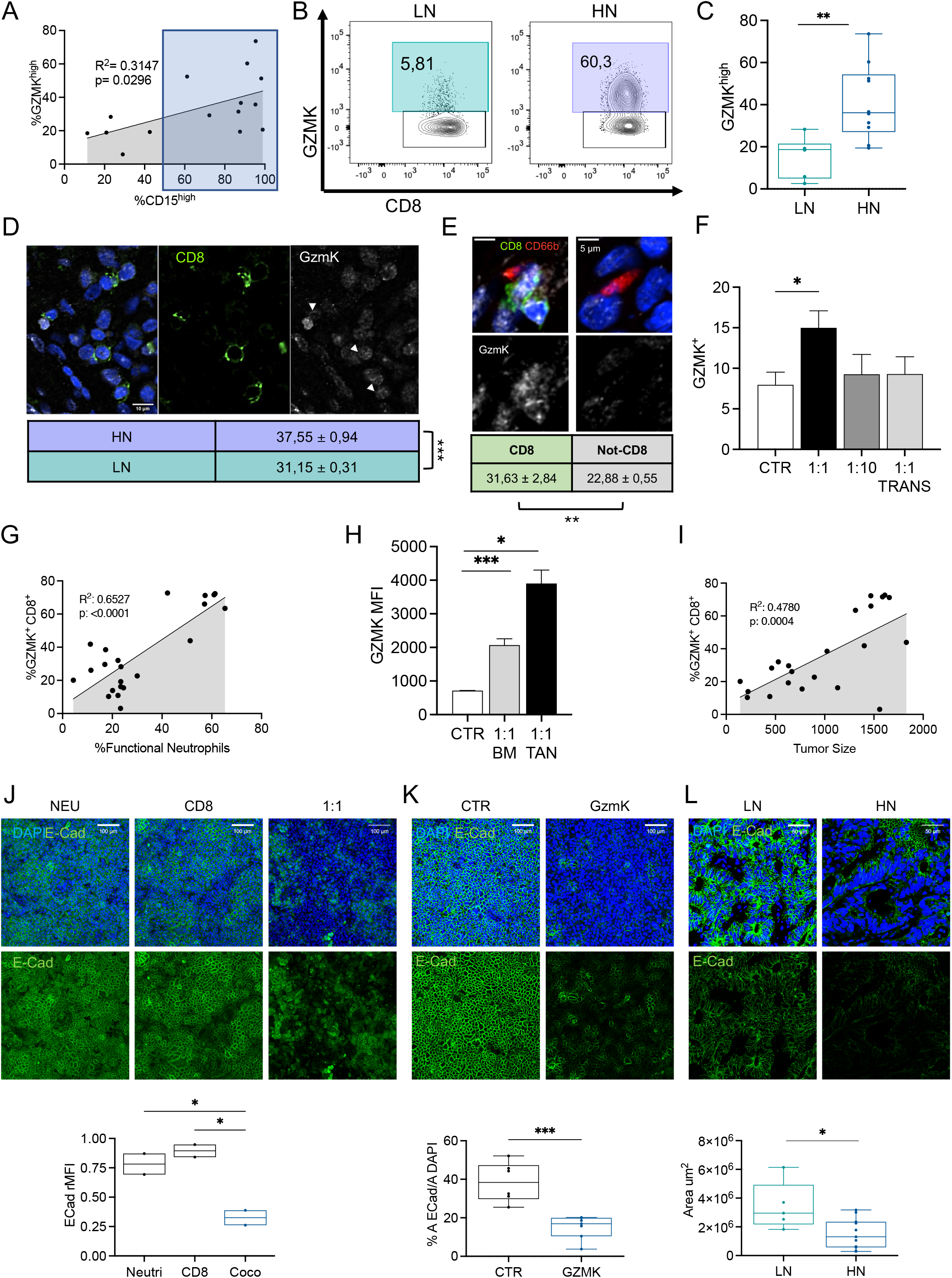
CD15^high^ neutrophil accumulation induces GZMK expression in CD8^+^ T cells, leading to tissue remodeling. **A.** Spearman correlation between CD15^high^ neutrophils and expression of GZMK within CD8^+^ T_EM_ cells on each individual CRC patient (black dots). Shadowed area (dark blue) includes High Neutrophils (HN) patients. **B.** Representative contour plots of CD8^+^ T_EM_ GZMK expression in Low Neutrophils (LN, turquoise) and High Neutrophils (HN, purple) patients. Percentage of CD15^high^ neutrophils are indicated. **C.** Box plot of GZMK^+^ CD8^+^ T_EM_ cells in HN (n=10) and LN (n=6) patients. **D-E**. Representative confocal images and quantification of GZMK Mean Fluorescence Intensity (MFI) in (**D**) HN (n=9) and LN (n=6) tumors. Scale bar 10mm. **(E)** CD8^+^ and CD8^−^ cells within a 10 μm distance from neutrophils (CD66b^+^). Scale bar 5 mm. (See Methods for details). **F.** Co-culture of CD8 and neutrophils isolated from peripheral blood of healthy donors. Bar plot of frequency of GZMK^+^ within CD8^+^ T cells after co-culture, in plate or in Transwell (TRANS) with neutrophils derived from healthy donors (HD). Neutrophils-to-CD8^+^ T cells ratio in the co-cultures are indicated. **G-I.** MC38 tumor bearing mice were euthanized and neutrophils content and GZMK expression in tumors evaluated by flow cytometry. **(G)** Spearman correlation between frequency of GZMK^+^ within CD8^+^ T cells and frequency of CXCR2^+^ cells within Ly6G^+^ CD11b^+^ neutrophils in MC38 tumors. **(H)** Bar plot of GZMK^+^ MFI within activated CD8^+^ T cells co-cultured in a 1:1 ratio with neutrophils isolated from Bone Marrow (BM) or tumors from MC38 tumor bearing mice. **(I)** Spearman correlation between frequency of GZMK^+^ within CD8^+^ T cells in MC38 tumors and tumor size (mm^3^). **J-K.** Representative confocal images and quantification of CACO-HT29 cells co-cultured on transwells and then (**J**) exposed for 24h to neutrophils (NEU), CD8^+^ T cells (CD8) or neutrophils/CD8^+^ T cells in 1:1 ratio (1:1), or (**K**) treated for 24h with control media (CTR) or active recombinant human GzmK (GzmK). Staining for DAPI in blue and E-Cadherin in green. Scale bar 100μm. **L.** Representatives confocal images and quantification of FFPE sections of LN and HN tumors from CRC patients. Staining for DAPI in blue and E-Cadherin in green. Scale bar 50μm.

Despite comparable frequencies of GZMK^high^ CD8^+^ T_EM_ cells in NAT and PB of LN and HN patients (Figure Supp. 4G), intratumor CD8^+^ T_EM_ cells expressed higher levels of GZMK in HN compared to LN patients, as assessed both by FACS (Figure 4B-C) and quantitative confocal microscopy (Figure 4D), confirming that the higher proportion of GZMK^high^ CD8^+^ T_EM_ cells was a tumor-specific hallmark in HN CRC patients.

Next, we investigated if neutrophils were directly involved in inducing a GZMK^high^ phenotype. In CRC patients, CD8^+^ T cells in proximity to neutrophils (distance r ≤10υm) expressed higher GZMK levels compared to other cell types (Figure 4E), suggesting that the interaction might be contact-mediated and the effect direct. Indeed, co-culturing of optimally activated CD8^+^ T cells and neutrophils derived from healthy donors demonstrated that the frequency of GZMK^high^ CD8^+^ T cells increased in presence of neutrophils and this was dependent from both the ratio of the two cell types and their physical contact (Figure 4F). These data revealed that skewing of CD8^+^ T cells toward a GZMK^high^ phenotype resulted from direct cross-talk between neutrophils and CD8^+^ T cells at the tumor site.

### GZMK^high^ CD8^+^ T_EM_ cells promote relapse in early CRC

To gain insight into the contribution of GZMK^high^ CD8^+^ T_EM_ and their interaction with neutrophils to the anti-tumor response in CRC, we analyzed the immune infiltrate during tumor progression in the syngeneic MC38 mouse model of colon cancer (Corbett et al., 1975). Similarly to what we observed on CRC patients, intratumor GZMK^high^ CD8^+^ T cells were positively correlated with the percentage of intratumor neutrophils (defined as CD11b^+^Ly6g^+^CXCR2^+^) in the MC38 model (Figure 4G). In addition, murine CD8^+^ T cells increased GZMK expression upon co-culturing with neutrophils from bone marrow and, to a larger extent, with tumor-infiltrating neutrophils (Figures 4H). Finally, the frequencies of intratumor GZMK^high^ CD8^+^ T cells and tumor-infiltrating neutrophils were positively correlated with the size of tumors (Figures 4I). Overall, these results confirmed the existence of a cross-talk between neutrophils and CD8^+^ T cells resulting in an increased GZMK production, and supported a model where GZMK^high^ CD8^+^ T cells might foster tumor progression.

Emerging evidence suggests that GZMK might promote inflammation and untargeted tissue damage (Wensink et al., 2015). This has been related to the activity of GZMK as an extracellular protease and its ability to promote epithelial-to-mesenchymal transition (EMT) by remodeling components of the extracellular matrix (Bouwman et al., 2021). We tested this hypothesis *in vitro* by exposing a functional human intestinal epithelial model to the neutrophils-CD8^+^ T cells co-culture, which we showed to induce GZMK production by CD8^+^ T cells. Compared to each of the two cell types alone, the co-culture induced a significant decrease in E-Cadherin expression (Figure 4J), a feature associated to EMT and tumor malignancy (MA et al., 2005; Zeisberg and Neilson, 2009). Similar results were obtained when the intestinal epithelial model was treated with purified GZMK (Figure 4K), providing evidence that release of GZMK would be sufficient to induce tissue remodeling. In line with this, HN CRC tumors showed reduced expression of E-Cadherin compared to LN (Figure 4L), suggesting that infiltration of GZMK^high^ CD8^+^ T cells could be linked to tumor progression and, possibly, contribute to relapse.

To test this link, we profiled patients in our cohort that encountered early relapse (Rel, n=5) and we observed that frequencies of CD15^high^ neutrophils and GZMK^high^ CD8^+^ T_EM_ cells in their T were comparable to HN and significantly higher than LN (Figure 5A-B). In agreement, T from relapsed patients were enriched in CL4 (mainly composed of CD8^+^ T_EM_ GZMK^high^CD39^neg^), while CL12 (mostly CD8^+^ T_EFF_ GZMK^low^ CD39^pos^) was poorly represented (Figures 1H-I and 5C-E). All together, these results support the existence of a CD8-neutrophil crosstalk at the tumor site, which significantly contributes to the clinical outcome (Pagès et al., 2009b, 2009c). Thus, evaluation of the different subtypes of cytotoxic CD8^+^ T cells could provide a mean to predict tumor recurrence and patient survival.

**Figure 5.**
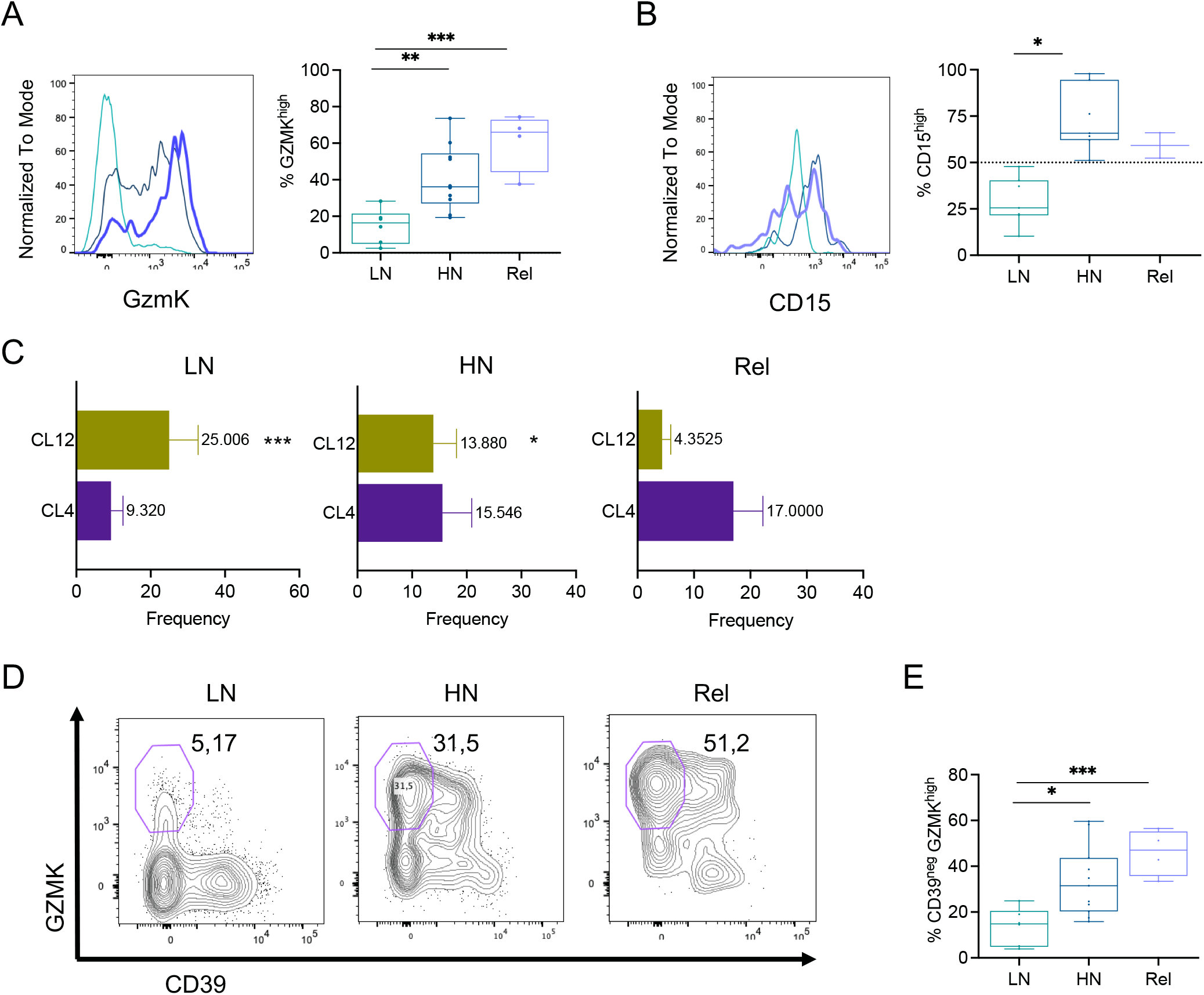
CRC patients encountering early relapse have high infiltration of GZMK^+^ CD8^+^ T effector memory (T_EM_) cells. **A.** Representative histogram and quantification of GZMK frequency within T_EM_ CD8 Tumor Infiltrating Lymphocytes (TILs) in High Neutrophils (HN), Low Neutrophils (LN) and Early-Relapsed (Rel) patients. **B.** Representative histogram and quantification of CD15^high^ neutrophils in HN, LN, Rel tumors. **C.** Bar plot of Cluster 12 (CL12, green) and Cluster 4 (CL4, purple) in HN, LN and Rel tumors. **D.** Representative contour plot of GZMK and CD39 expression within CD8^+^ T in in HN, LN and Rel tumors. **E.** Box plot of CD39^−^ GZMK^+^ within CD8^+^ T cells in HN, LN and Rel tumors.

### GZMK expression correlates with a distinct transcriptional program in CD8^+^ T_EM_

Aiming to build a signature that could address the prognostic potential of infiltrating GZMK^high^CD8^+^ T_EM_ cells, paired T and NAT tissues from 8 CRC (stage I-III) HN patients were profiled by single cell RNA sequencing (scRNAseq). We collected data from a total of 52316 cells. Mean reads per cell were 94183 (min:37500; max:235694) with a median number of genes detected per cell of 3779 (min:1433; max:5392). Sequencing saturation was >49% for all samples, indicating comprehensive sampling of the available transcripts. After processing and normalization, dimensionality reduction was performed with uniform manifold approximation and projection (UMAP). Shared nearest neighbor (SNN) clustering distinguished eleven different clusters that were annotated to nine broad cell lineages, including epithelial (*EPCAM^+^*), endothelial (*CD34^+^*), mesenchymal (*MMP2^+^*), B cells (*MZB1^+^*), and T/NK (*CD3D^+^*) (Figures Supp. 5A-C). Notwithstanding the expected variability among patients, all the annotated clusters were represented across the patients (Figure Supp. 5D). Expression analysis across all the annotated cell subtypes highlighted the T/NK cluster as the main source of *GZMK* expression in the TME of HN (Figure Supp. 5E).

Zooming in on the 8523 cells composing the T/NK cluster, we generated a new UMAP that further distinguished seven T cell subtypes (Figure Supp. 5F), among which cluster 1 was enriched in CD4^+^ while clusters 3 and 4 were enriched in CD8^+^ T cells (Figures Supp. 5G-H). Interestingly, *GZMK* expressing cells were mostly restricted to the CD8^+^ clusters 3 and 4, confirming results from our phenotypic analysis. Within the CD8^+^-enriched T cells clusters 3 and 4, we could identify eleven sub-clusters (Figures 6A). These clusters were annotated with manually curated gene-sets (Table Supp. 1) based on recent literature (Li et al., 2019). While confirming the presence of previously described putative T cell sub-populations, this analysis highlighted a remarkable transcriptional heterogeneity. The eleven clusters included minor fractions of CD4^+^ T cells falling in the memory (CD4_T_EM_), the *FOXP3* expressing regulatory (Treg), *CD40LG* positive mucosal associated invariant T (MAIT) and exhausted (Tex) T cell clusters. Co-expression of *TRDC* and *TRGC1* enabled discrimination of gamma delta T cells (Tγδ), while higher expression of *AKR1C3*, *CXC3R1*, *FCER1G*, *NKG7* and *PRF1* were distinctive transcriptional traits of the natural killer (NK) cluster. Naive-like T cells were assigned based on expression of *IL7R*, *CCR7* and *SELL* signature genes (Figures Supp. 6A and C). Most of the retrieved T cells fell within the CD8^+^ T effector memory subgroup (CD8_T_EM_, Figure Supp. 6B) characterized by low expression of *GZMB* and high expression of *GZMK* (Figures 6B), in agreement with the GZMK^high^ T cells identified by FACS analysis. On the other hand, marginal expression of genes associated to exhaustion, such as *CTLA-4*, *PDCD-1*, *CXCL13*, *HAVCR2*, *ENTPD1* and *LAYN*, and differential expression analysis revealed that CD8_T_EM_ were considerably distinct from T_EX_ cells (Figures 6C-D). In addition, pseudotime analysis showed that CD8_T_EM_ were closer to Naive-like T cells than to terminally differentiated effectors (T_TE_, Figures 6E and Supp. 6D). The borders between CD8_T_EM_ and the clusters labeled as CD8_T_EM_-PD1 and HLADR_CD8 were diffuse, instead, even though hierarchical clustering of differentially expressed genes confirmed the robustness of the model. While suggesting that transcriptional gradients may contribute to T cell heterogeneity, this observation also marks intermediate states on the transition of CD8_T_EM_ toward exhaustion, for example, through the expression of genes like *PDCD1* and *CXCL13*. Functionally, pathway enrichment analysis indicated that, despite having lower cytolytic potential (padj= 0.005) compared to T_TE_ and NK cells, both CD8_T_EM_ and HLADR_CD8 (both expressing high levels of *GZMK* levels) were not impaired in proliferation (Figure Supp. 6E). In particular, CD8_T_EM_ had lower expression of lymphocyte activation and migration and higher cell-cell adhesion signatures (Figure Supp. 6E). Noteworthy, similar expression levels of effector molecule *TNFα* characterized the two subsets of CD8_T_EM_ grouped based on *GZMK* expression (Figures Supp. 6F-G), further supporting that *GZMK* expression is not associated with a dysfunctional state in CD8_T_EM_ (see also Figure Supp. 3D). Finally, querying of publicly available CRC datasets on The Cancer Genome Atlas (TCGA) using a transcriptional signature that distinguished CD8_T_EM_ highly expressing *GZMK* among those defined by scRNAseq (Figure Supp. 6I, see Methods for details), showed better prognosis in patients with lower infiltration of CD8 T_EM_ expressing high *GZMK* (Figure 6G). This signature provided a better prognostic value than the functional signature related to the cytolytic activity of CD8^+^ T cells and it was effective in stratifying patients with other carcinoma types, like the TCGA lung adenocarcinoma (Figure Supp. 6J).

**Figure 6.**
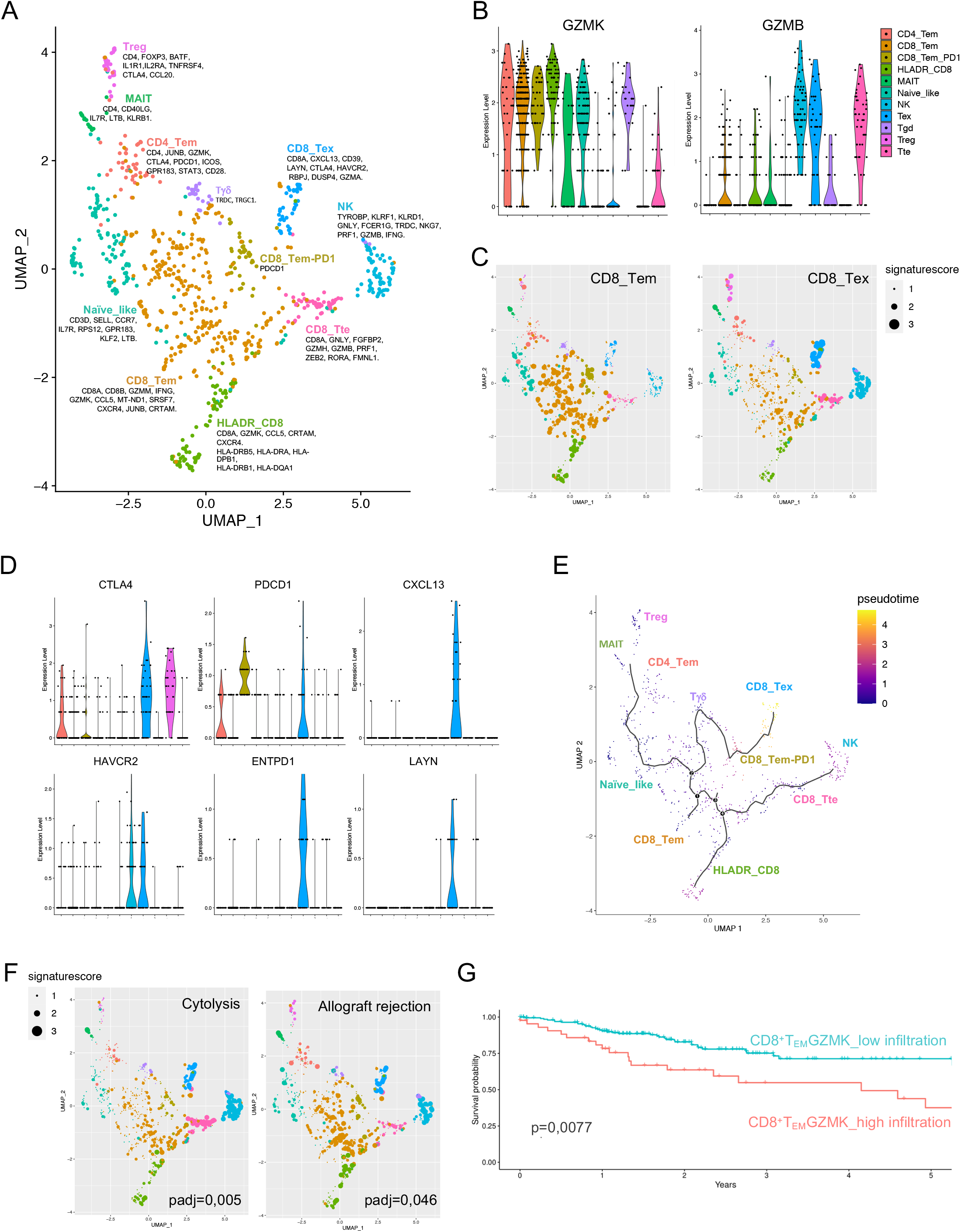
Single-cell RNAseq analysis on CRC patients defines an effective prognostic signature based on CD8^+^ GZMK^+^ T_EM_ abundance. UMAP projection of CD8^+^ T cells. Colors represent different T cell subtypes. Key genes used for manual annotation are indicated. **B.** Violin plots showing the normalized expression of GZMB and GZMK in the different T CD8^+^ cell subtypes. **C.** UMAP showing the expression of CD8_Tem and CD8_Tex signatures for each cell. Cells are color coded based on SNN-clustering, dot size reflects the expression of the signature for each cell. **D.** Violin plots showing the expression of genes associated with T-cell exhaustion within the indicated cell subtype. **E.** Monocle pseudotime analysis of the CD8^+^ T cell subtypes. The pseudotime line connects the indicated T cell subtypes identified in (A). **F.** UMAP as in (C) showing the expression of Cytolysis and Allograft rejection signatures. **G.** Kaplan-Meyer analysis of the association of GZMK^high^ cell subtype’ abundance with disease free survival (DFS) on the TCGA-COAD cohort (see Methods and Table Suppl. 1 for details). High signature in red, low in turquoise.

In conclusion, our functional and transcriptional characterization of the tumor immune infiltrates resulted in an effective stratification of patients and predicted a different risk of relapse in early-stage CRC patients. Our findings have the potential to be extended to other cancer types, including lung carcinoma.

## DISCUSSION

We conducted unbiased analysis of the immune cells within and across CRC patients by combining multiparametric flow cytometry with functional assays and single cell transcriptomics. Our data reveal that CRC patients can be stratified based on high (HN) or low (LN) CD15^high^ neutrophil infiltration and highlight the emergence of a novel GZMK^high^ CD8^+^ T_EM_ cell population as a hallmark of HN tumors (Figure 7). The predominance of the GZMK^high^ CD8^+^ T_EM_ cells in CRC patients that encountered an early relapse and the correlation with poor clinical outcome on an independent cohort underline the clinical relevance of the cell population and its therapeutic potential.

**Figure 7.**
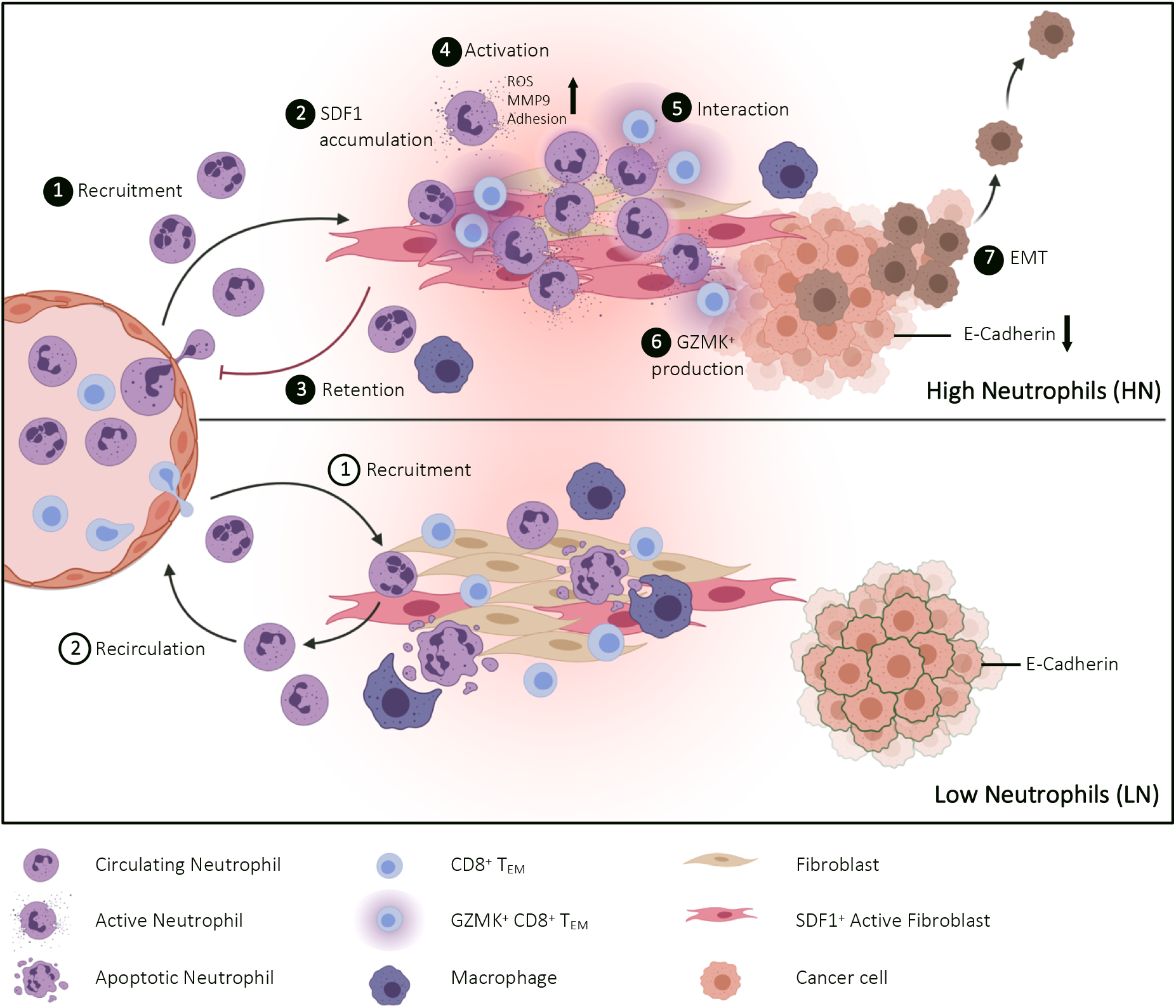
Simplified graphical summary. Abundance of CD15^high^ neutrophils in the tumor identifies High (HN) and Low (LN) neutrophil subgroups of early-stage CRC patients (**1**). Elevated SDF-1 levels in HN tumors reshape the functional state of infiltrating neutrophils (**2**), promoting their retention (**3**) and activation (**4**) at the tumor site. The interaction with neutrophils (**5**) pushes effector memory CD8^+^ T (T_EM_) to produce high levels of Granzyme K (GZMK) (**6**), which in turn remodels the tumor microenvironment (TME) fostering EMT (**7**). A GZMK^high^ T_EM_ transcriptional signature effectively stratify non-metastatic early CRC patients and predict poor clinical outcome.

Despite several attempts to profile the immune cell content of CRC (Pagès et al., 2009b; Zhang et al., 2018), the prognostic value of immune cell infiltration is not conclusive, especially regarding neutrophils (Shaul and Fridlender, 2019). We provide evidence that neutrophils are the most abundant innate cell population infiltrating the tumor bed in early-stage CRC. Pro-tumorigenic roles have been ascribed to neutrophils across different tumor types (Galdiero et al., 2018; Mantovani et al., 2011), including promotion of angiogenesis and immunosuppression (Bronte et al., 2016; Coffelt et al., 2016), but it is not clear whether distinct neutrophil populations gather the tumor or if phenotypic changes occur inside the TME. The presented analysis of the canonical neutrophil marker CD15 (Pillay et al., 2013) reveals phenotypic changes specifically in the population associated with tumors. Intratumor neutrophils from LN patients have significantly lower expression of CD15 compared to healthy- and patient-derived peripheral neutrophils. Conversely, neutrophils infiltrating the TME of HN patients maintain high levels of CD15 expression, are enriched in CXCR2^+^CXCR4^+^ cells and tend to produce more ROS, in line with a so called N2-like tumor-promoting functional state (Fridlender et al., 2009). Interestingly, treatment of freshly isolated neutrophils with interstitial fluid from HN tumors led to higher CD15 expression but, differently from interstitial fluid from LN, did not foster migration, suggesting that specific soluble factors in the TME might polarize neutrophils toward a CD15^high^ state rather than increasing their recruitment. Neutrophils are recruited early to the site of inflammation, where their not-specific antimicrobial functions can endanger the integrity of the host tissue in case of inappropriate retention (Nathan and Ding, 2010; Rock et al., 2010). To avoid it, in addition to clearance by macrophages-directed efferocytosis (Cox et al., 1995; Grigg et al., 1991), neutrophils redistribute into surrounding tissues or vasculature through reverse migration (Buckley et al., 2006; Robertson et al., 2014; Wang et al., 2017; Woodfin et al., 2011), which has been suggested to follow desensitization to local chemotactic gradients (Holmes et al., 2012; Powell et al., 2017; Robertson et al., 2014). In this regard, SDF-1/CXCR4 signaling plays an important role in the retention of neutrophils at inflammatory sites (Isles et al., 2019). Since clearance by resident tissue macrophages has been associated with reduction of CD15 surface expression on neutrophils (Shelley et al., 2018), we can speculate that the enrichment of CD15^high^ neutrophils in HN tumors might be due, at least in part, to inefficient clearance, as previously demonstrated in other tumor contexts (Peng et al., 2021a). On the other hand, our analysis of differentially abundant soluble factors pointed at SDF-1, which correlates *in vivo* with frequency of CD15^high^ neutrophils and, *in vitro,* increased adhesion and release of TME’s remodeling factors rather than driving chemotaxis. Overall, these data suggest that elevated concentrations of SDF-1 may promote the reprogramming of neutrophils infiltrating HN tumors. Future studies would need to address if this may lead to an increased retention of neutrophils in the tumor, altering their homeostatic turnover.

Neutrophils influence CD8^+^ T cell-mediated responses both in animal models and humans. However, results are still sparse and often conflicting (Eruslanov et al., 2014; Governa et al., 2017; Jaillon et al., 2020b; Li et al., 2020; Sangaletti et al., 2014b; Singhal et al., 2016; Wu et al., 2014). We provide evidence that direct interaction between neutrophils and CD8^+^ T cells leads to increased GZMK expression, both *in vivo* and *in vitro*. When compared to their GZMK^neg^ counterpart, GZMK^high^ CD8^+^ T_EM_ cells harbor significantly lower PD-1 expression and higher levels of Gzmβ, posing these cells as distinct compared to the GZMK^+^ CD8^+^ T cell subpopulations previously identified and described as pre-dysfunctional (Guo et al., 2018; Hornburg et al., 2021; Mogilenko et al., 2021; Zhang et al., 2018). Moreover, their abundance in the TME is independent from tumor stage and aging (P > 0.05, ANOVA) (Mogilenko et al., 2021). Of particular relevance, GZMK^high^ CD8^+^ T_EM_ cells lack CD39 expression, which might suggest that they are contributing to a general inflammatory state (Chiou et al., 2021; Corridoni et al., 2020) rather than targeting specifically tumor associated antigens (Scheper et al., 2019; Simoni et al., 2018). Thus, the high proportion of GZMK^high^ CD39^neg^ CD8^+^ T_EM_ cells found in HN CRC patients raises the possibility that they could be involved in a cross-reactive response, favoring cancer progression. The alloreactive-like transcriptional signature associated with GZMK^high^ CD8^+^ T_EM_ favors this model, with upregulation of CRTAM and CCL5/RANTES promoting retention of activated GZMK^high^ CD8^+^ T_EM_ cells (Arase et al., 2005; Cortez et al., 2014; Galibert et al., 2005) and recruitment and immunosuppressive polarization of myeloid cells (Aldinucci et al., 2020; Schall et al., 1990; Seo et al., 2020; Swanson et al., 2002; Walzer et al., 2003), respectively. However, only future analysis of the TCR clonality of GZMK^high^ CD39^neg^ CD8^+^ T_EM_ will allow testing this hypothesis unequivocally.

We show that, at least in HN CRC patients, CD8^+^ T_EM_ are the main source of GZMK. Further studies are needed to determine whether CD8^+^ T cells recruited to the tumor are skewed by neutrophils toward a GZMK^high^ phenotype or, conversely, the presence of GZMK^high^ CD8^+^ T cells impacts neutrophil homeostasis and function, as already proved for other immune cell populations (Joeckel et al., 2011). Regardless of the order of events, GZMK might play a pathogenic role by contributing to inflammation and tissue damage. This relates to its non-cytotoxic activity as extracellular protease and its ability to promote cellular migration and epithelial-to-mesenchymal transition (EMT) by remodeling components of the extracellular matrix (Joeckel and Bird, 2014; Wensink et al., 2015), facilitating cytokine responses (M et al., 2016; Wensink et al., 2014) In line with this, HN tumors showed reduced expression of E-Cadherin compared to LN, which is a feature of EMT and tumor malignancy. Along with tangible therapeutic implications for future interventions, high expression of GZMK by infiltrating CD8^+^ T_EM_ in response to CD15^high^ neutrophils links them to tumor progression rather than anti-tumor function, suggesting that integrating functional information resulting from the crosstalk of CD8^+^ T cells with different components of the immune contexture and the stroma might implement the prognostic value of current biomarkers, which are mostly based on limited phenotypic or transcriptional CD8^+^ T cell characteristics.

## LIST OF ABBREVIATIONS

CRC: Colorectal Cancer
GZMK: Granzyme K
CXCL12/SDF-1: Stromal Cell-Derived Factor 1
TNM: Tumor-Node-Metastasis
TME: Tumor Microenvironment
Tils: Tumor Infiltrating Lymphocytes
PB: Peripheral Blood
T: Tumor
NAT: Normal Adjacent Tissue
DCs: Dendritic Cells
HN: High CD15^high^ Neutrophils
LN: Low CD15^high^ Neutrophils
TRos: Total Reactive Oxygen Species
T_th_: CD4^+^ Th Cells
T_reg_: T Regulatory Cells
Tγδ: T Gamma-Delta Cells
T_N_: Naive T Cells
T_CM_: Central Memory T Cells
T_EM_: Effector Memory T Cells
T_EMRA_: Terminally Differentiated Effector Memory T Cells Re**-**Expressing CD45RA
Tsne: T-Distributed Stochastic Neighbor Embedding
MFI: Mean Fluorescence Intensity
CL: Clusters
T_RM_: Tissue-Resident Memory T Cells
MAIT: Mucosal Associated Invariant T Cells
EMT: Epithelial-To-Mesenchymal Transition
Rel: Early Relapse
scRNAseq: Single Cell RNA Sequencing
UMAP: Uniform Manifold Approximation And Projection
SNN: Shared Nearest Neighbor
T_ex_: Exhausted T Cells
NK: Natural Killer Cells
T_TE_: Terminally Differentiated Effectors Cells
TCGA: The Cancer Genome Atlas
PBMCs: Peripheral Blood Mononuclear Cells
PMNs: Polymorphonuclear Leukocytes
TEER: Trans-Epithelial Electrical Resistance CACO2
HMEC1: Human Micro-Endothelial Cells
Imfi: Integrated MFI
IHC: Immunohistochemistry
FFPE: Formalin-Fixed Paraffin Embedded
UMI: Unique Molecular Identifiers
SCMCA: Single Cell Mouse Cell Atlas
OS: Overall Survival
DFS: Disease-Free Survival
COAD: Colon Adenocarcinoma Cohort

## SUPPLEMENTARY INFORMATIONS

### Availability of supporting data

All data supporting the findings of this study are available from the corresponding authors upon reasonable request.

### Authors’ contributions

LN and TM contribute equally. Correspondence to Teresa Manzo. ST, TM and LN designed. ST, CC, CS, MB, CBNL, GR performed experiments; ST, CC, CS, OC, MB, DC, CS, CBNL, ADM, GB, VL, PP, GR, MS, SC, LN and TM analyzed data; MR, EL provided critical expertise and resources; WP, SPR, NF and UFR obtained tissues from patients; LN and TM coordinated the whole study; ST, CC, LN and TM wrote and edited the manuscript. All the authors discussed and read the manuscript.

## Acknowledgements

LN and TM dedicate this work to Leonardo and Lara Nezi. We thank Phil Greenberg for thoughtful discussion. The authors thanks IEO Biobank B4Me, all the nurses, the Molecular Pathology, Imaging and Genomic Unit at the Department of Experimental Oncology at IEO, patients abd their families involved in the study.

