## Supplementary Information for "GZMK^high^ CD8^+^ T effector memory cells are associated with CD15^high^ neutrophil abundance in early-stage colorectal tumors and predict poor clinical outcome"

### SUPPLEMENTARY FIGURE LEGENDS.

**Supplementary Figure 1 (related to Figure 1). Characterization of CD15$^{high}$ neutrophils infiltrating early-stage CRC. A.** Bar plot of dendritic cells (DC), macrophages and NK cells within CD45$^{+}$ in T and NAT tissue. **B.** Representative images and quantification of immunohistochemistry (IHC) of CD66b$^{+}$ cells in T and NAT tissues. Scale bar 100μm. **C.** Flow cytometric gating strategy for the identification of CD15$^{high}$ tumor associated neutrophils and representative dot plot in Low (LN) and High (HN) neutrophils patients respectively. **D.** Quantification of CD15 MFI within CD15high and CD15$^{low}$ neutrophils. **E.** Representative histogram of CD15 expression in neutrophils and bar plot quantification of CD15$^{high}$ neutrophils in T and NAT tissue within LN and HN patients. **F-G.** Frequency of (**F**) CD10$^{+}$ cells and (**G**) CXCR4$^{+}$ CXCR2$^{-}$ cells within CD15$^{low}$ and CD15$^{high}$ neutrophils in CRC tumors. **H-I.** Bar plot representation of (**H**) CXCR4$^{+}$ CXCR2$^{-}$ cells and (**I**) total ROS (tRos) MFI within neutrophils in LN and HN tumors from CRC patients. **J.** Spearman correlation between frequency of CD15$^{low}$ neutrophils and frequency of macrophages within CD45$^{+}$ cells in CRC tumors. **K.** Frequency of macrophages in LN and HN patients. Bars indicate 90-10 percentile. **L.** Fold change of CD15 expression on neutrophils from healthy donors (HD) treated with Interstitial Fluid from LN and HN patients. **M.** Representative confocal images and quantification of Stromal Derived Factor 1 (SDF-1) and αSMA in formalin-fixed paraffin embedded (FFPE) tumor tissues from LN and HN patients. DAPI in blue, αSMA in green, SDF-1 in red. See Methods for details. **N.** Representative confocal images and quantification of Stromal Derived Factor 1 (SDF-1) per αSMA$^{+}$ cells in LN and HN patients. DAPI in blue, αSMA in green, SDF-1 in red. **O-P.** Representative images and quantification of migrating neutrophils on a microfluidic chip upon stimulation with CXCL6 (400 ng/mL), IL-8 (100 ng/ml), SDF-1 (100 ng/ml) or culture medium. Scale bar 100μm. **Q-S.** Bar plot representation of the expression (MFI) of CXCR4, CXCR2 and CD62L on SDF-1 treated-neutrophils from HD. *, P < 0.05**, P < 0.01 ***, P < 0.001.

**Figure Supplementary 2 (related to Figure 2). High-dimensional single cell analysis of CD8$^+$ T cells.** Kaplan-Meyer analysis of the association of CD8 cytotoxic signature with disease free survival (DFS) on the TCGA-COAD cohort (see Methods, for details). High signature in red, low in turquoise. Table showing number of patients at risk in each group at indicated time points. **B.** Frequency of CD3$^+$ and CD3$^-$ cells within CD45$^+$ total immune infiltrate in T tissues. **C.** Representative dot plot of CD45RA and CCR7 expression within CD8$^+$ T cells in Peripheral Blood (PB), Normal Adjacent Tissue (NAT), tumor (T) tissue. **D**. Representative dot plot of CD28 and CD27 expression within CD8$^+$ T$_{EM}$. **E.** Eulero venn diagram of checkpoint inhibitor frequencies within CD8$^+$ TILs (n=31). **F.** Expression level of selected markers in the CD8$^+$ T cells tSNE representation of concatenated PB, NAT and T samples. **G.** Balloon plot of average expression levels and frequencies of 18 selected markers in the analyzed CD8$^+$ T cells clusters (CL). Hierarchical clustering based on markers and CL. **H.** Unsupervised clustering analysis of mean frequencies of 20 CL identified. ***; P < 0.001.

**Figure Supplementary 3 (related to Figure 3). Phenotypic characterization of CD39$^-$ GZMK$^+$ CD8$^+$ T cells in the blood and tumor of CRC patients. A-B.** Representative histogram and quantification of (**A**) CD69 and (**B**) CD103 frequency within CD39$^-$ CD8$^+$ T$_{EM}$ in Tumor (T) and Peripheral blood (PB). **C-D.** Bar plot of (**C**) GZMK$^+$ and (**D)** GzmB$^+$, IFNγ$^+$ and TNFα$^+$ frequency within sorted CD39$^+$ and CD39$^-$ CD8$^+$ T$_{EM}$ from T tissues. *, P < 0.05 ;**, P < 0.01 ***; P < 0.001.

**Figure Supplementary 4 (related to Figure 4).** Kaplan-Meyer analysis of the association of combined neutrophil infiltration (Neutr) and $CD8^{+}$ cytolytic (Cyt) signatures with disease free survival (DFS) on the TCGA-COAD cohort (see Methods, for details). No Neutr and High Cyt in red, No Neutr and Low Cyt in green, Yes Neutr and High Cyt in turquoise and Yes Neutr and Low Cyt in purple. Table showing number of patients at risk in each group at indicated time points. **B.** Frequency of naïve ($T_{NAIVE}$), Central Memory ($T_{CM}$), Effector Memory ($T_{EM}$) and Effector Memory $CD45RA^{+}$ ($T_{EMRA}$) T cells in High Neutrophils (HN, blue) and Low Neutrophils (LN, dark green) patients. **C.** Frequency of indicated markers within $CD8^{+}$ $T_{EM}$ cells in HN and LN patients. **D.** Frequency of $GZNK^{+}$ cells within $CD4^{+}$ $T_{EM}$ cells. **E.** Spearman correlation between $CD15^{high}$ neutrophils and $GZMK^{+}$ $CD4^{+}$ $T_{EM}$ cells. **F.** Representative contour plot of GZMK expression in $CD4^{+}$ $T_{EM}$ cells, and quantification of $GZMK^{+}$ $CD4^{+}$ $T_{EM}$ cells cells in LN and HN patient. swithin $CD4^{+}$ $T_{EM}$ cells. **G.** Frequency of $GZMK^{+}$ $CD8^{+}$ $T_{EM}$ cells in Tumor (T), Normal Adjacent Tissue (NAT) and Peripheral Blood (PB) of LN and HN patients**, $P < 0.01$.

**Figure Supplementary 5. Single-cell RNAseq analysis of the immune cell compartment infiltrating HN CRC patients.**

**A.** UMAP projection of cells of the 8 patients considered for the analysis. Cells were colored based on the SNN-clustering. Each cluster was assigned to a cell type, as indicated (see Methods). **B.** Heatmap showing the scaled UMI counts of the differentially expressed genes found in each cluster. **C.** Violin plots showing the normalized expression of select cell type markers in the SNN cluster shown in (A). **D.** Bar plot showing patients' distribution of each cell cluster. Colors match the indicated tissue of origin. **E-F.** UMAP projection of cells belonging to the T/NK cluster shown in (A). (E) Colors represent the SNN clusters. (F) The expression of indicated T cell markers is shown. **G.** Violin plots showing the expression of selected genes for each cluster in T/NK cells.

**Figure Supplementary 6. Single-cell RNAseq analysis of the CD8$^+$ T cell compartment infiltrating HN CRC patients.**

**A.** Bubble plot showing the expression of selected markers for each manually-annotated T cell subtype (see Table Suppl. 1 for list of genes used in the annotation). The size of the bubble represents the fraction of cells with at least one UMI for a specific gene, while the color shows the median of the scaled normalized expression in that fraction of cells. **B.** Bar plot showing the number of cells in each CD8$^+$ T cell subtype. **C.** Violin plots showing the expression of select marker genes in each of the indicated T cell subtypes. **D-E.** UMAP showing the expression of genes selectively expressed in (D) each cell type (see Suppl. Table 1) or (E) immune signature (Reactome). Cells are color coded based on SNN-clustering; dot size reflects the expression of the signature in each cell. **F.** Distribution plot showing GZMK expression in the indicated CD8$^+$ T cell subtypes. **G.** Violin plots showing the expression of GZMK and TNF$\alpha$ in the entire CD8$^+$ T cell population and in CD8$^+$ $T_{EM}$ cells. Cells were considered positive for GZMK if the number of GZMK UMI was ≥2. **H.** Bubbleplot showing the expression of differentially expressed genes between GZMK$^+$ and GZMK$^-$ CD8$^+$ Tem cells. The size of the bubble represents the fraction of cells expressing a specific gene (n UMI ≥1), while the color shows the median of scaled normalized expression of that fraction of cells. The threshold for considering a cell as GZMK$^+$ was set to the number was set to the number of UMI≥2. **I.** Distribution of GZMK$^{high}$ cell subtype' infiltration across TCGA-COAD samples. The optimal cutpoint for stratifying high and low level GZMK$^{high}$ cell subtype' infiltration was obtained by using the Maximally Selected Rank Statistics. **J.**

Kaplan-Meyer analysis of the association of GZMK$^{high}$ cell subtype' abundance with overall survival (OS) on the TCGA-LUAD cohort (see Methods and Table Suppl. 1 for details).

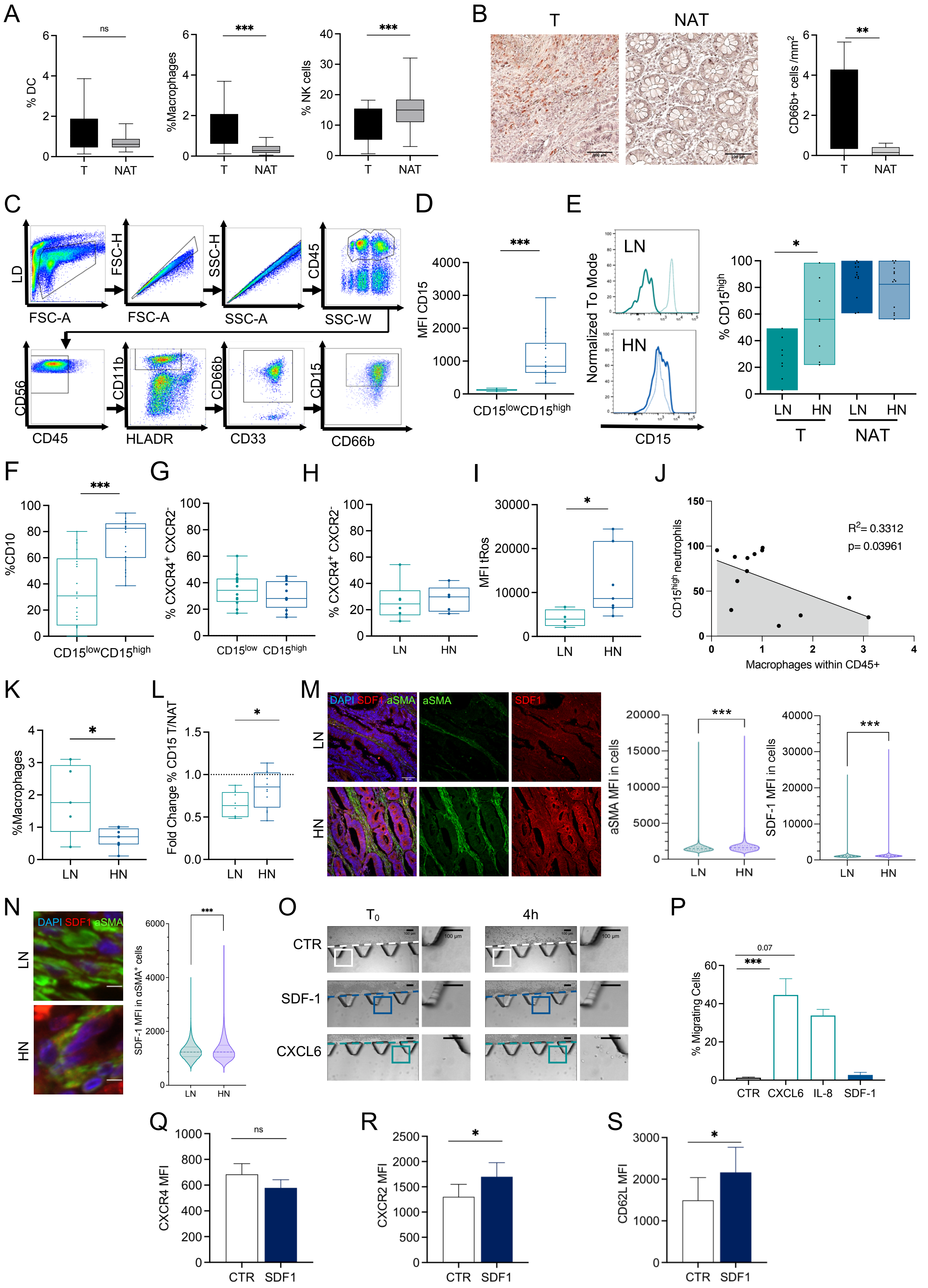


**Figure Supp. 1**

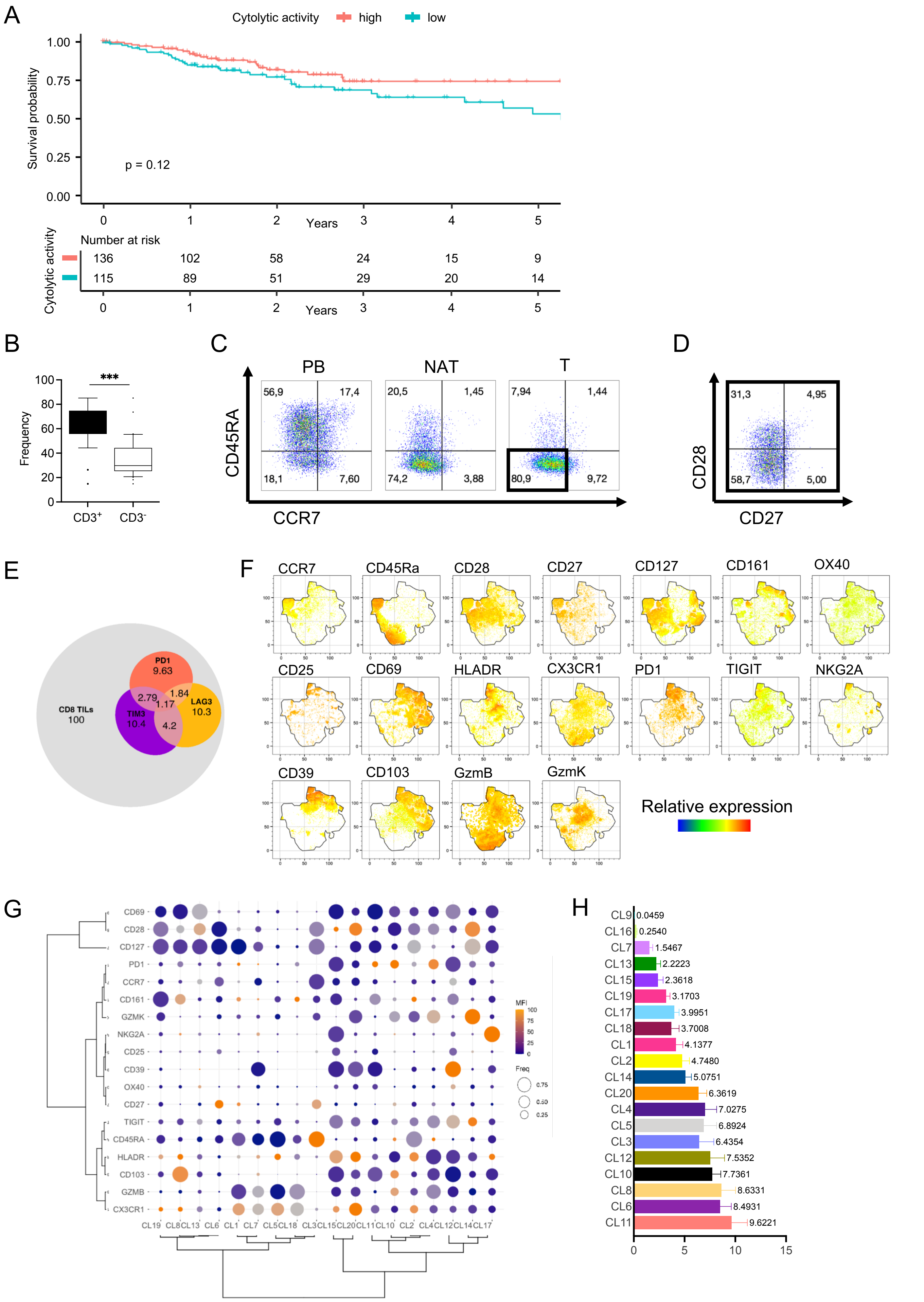


**Figure Supp. 2**

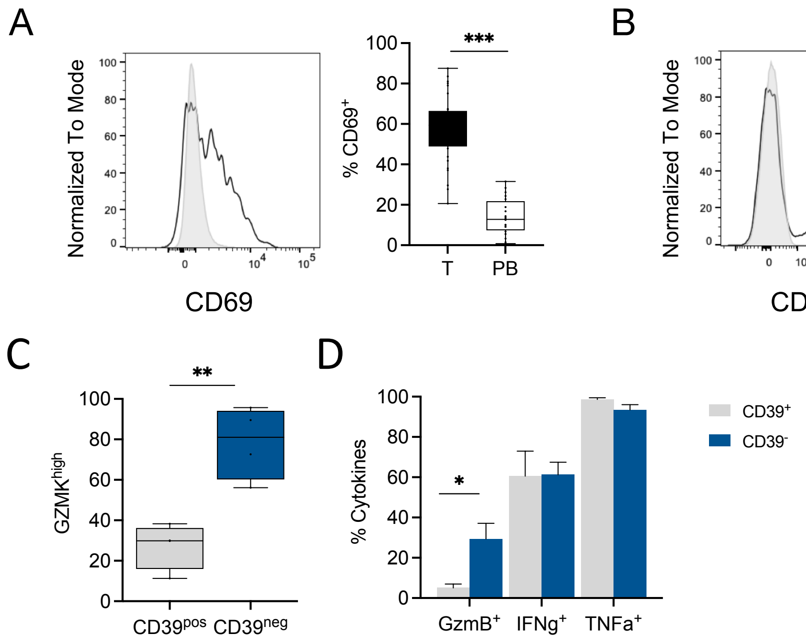


**Suppl. Figure 3**

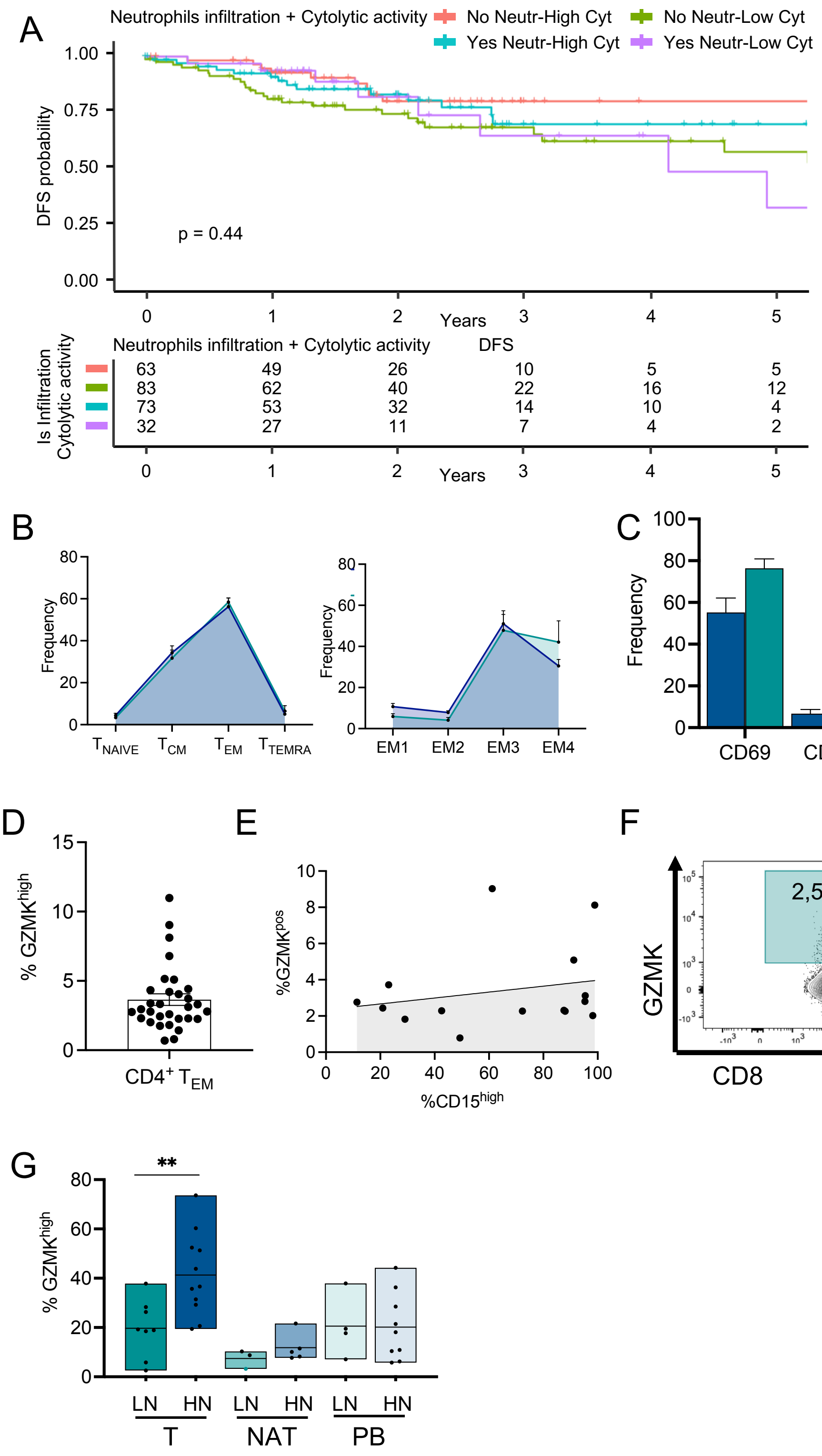


**Suppl. Figure 4**

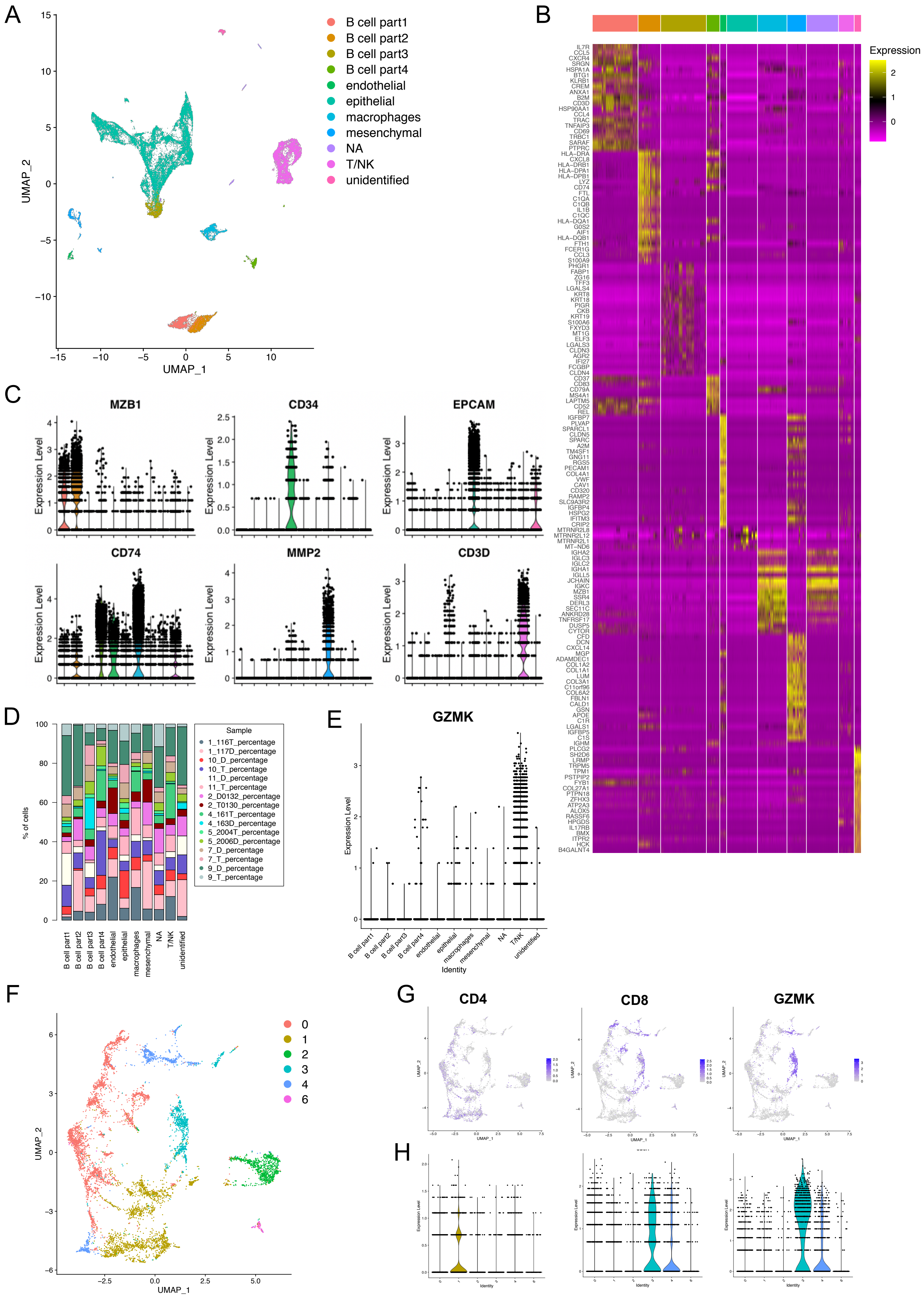


**Figure Supp. 5**

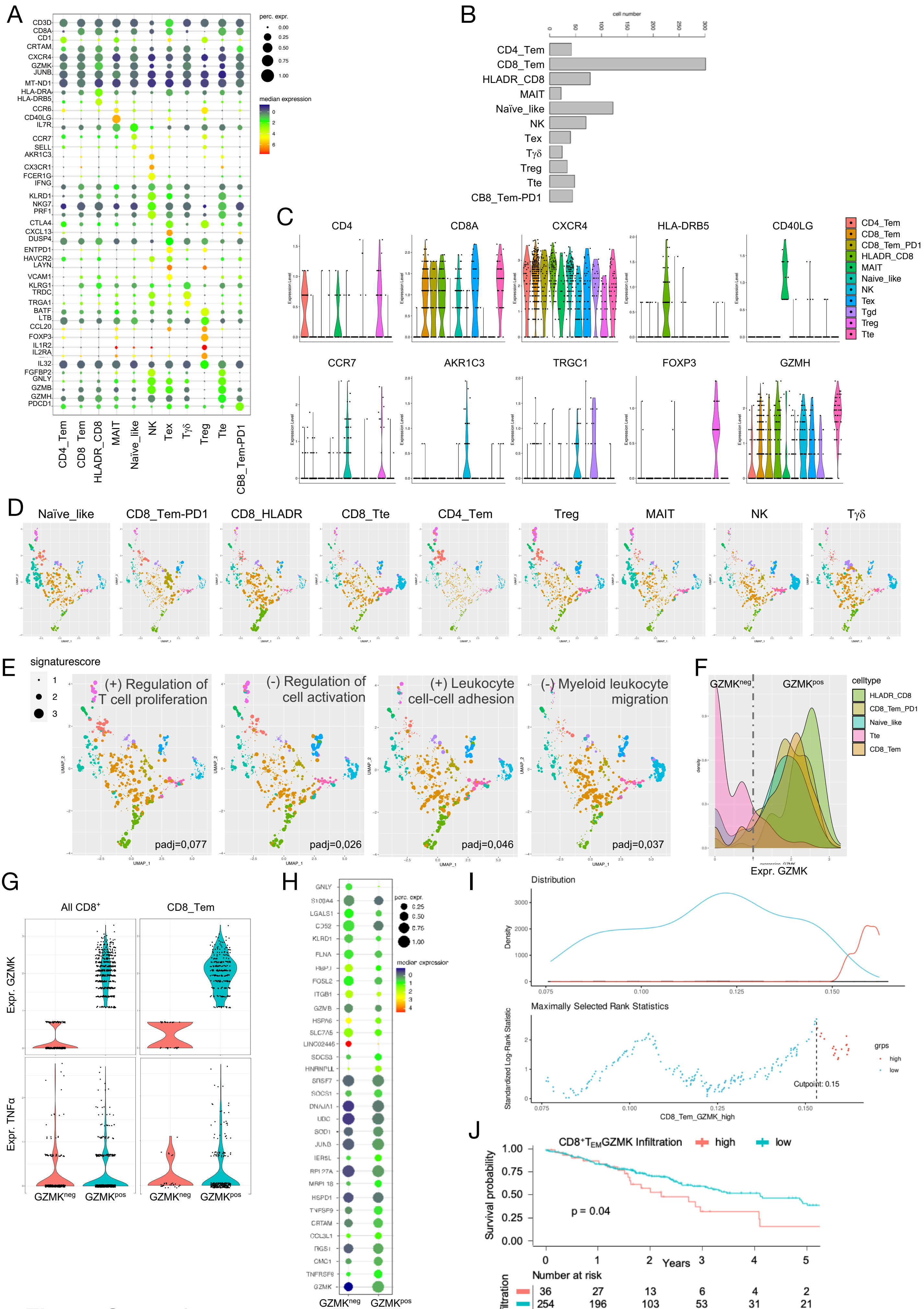


**Figure Supp. 6**

| T cell population | Gene Signature |
|---|---|
| CD8 Effector memory T cells (CD8_Tem) | CD3D, CD3E, CD8A, CD8B, GZMM, IFNG, GZMK, CCL5, MT-ND1, SRSF7, CXCR4, JUNB, CRTAM. |
| CD8 Effector memory T cells (CD8_Tem-GZMK$^{high}$) | CD3D, CD3E, CD8A, CD8B, IFNG, GZMK, CCL5, MT-ND1, SRSF7, CXCR4, JUNB, CRTAM, TNFRSF9, CMC1, JUN, CCL3L1. |
| CD8 Effector memory T cells-HLADR CD8 (CD8_Tem-HLADR) | CD3D, CD3E, CD8A, CD8B, CCL5, JUNB, CRTAM, HLA-DRB5, HLA-DRA, HLA-DPB1, HLA-DRB1, GZMK, HLA-DQA1, CD74, ZFP36, CXCR4. |
| CD8 Effector memory T cells PD-1 (CD8_Tem-PD1) | CD3D, CD3E, CD8A, CD8B, GZMK, CCL5, MT-ND1, CXCR4, CRTAM, PDCD1. |
| CD8 Terminal effector T cells (CD8_Tte) | CD3D, CD3E, CD8A, CD8B, GNLY, FGFBP2, FCGR3A, GZMH, GZMB, TRGC2, PRF1, ZEB2, RORA, MT-ND3, FMNL1. |
| CD8 Exhausted T cells (CD8_Tex) | CD3D, CD3E, CD8A, CD8B, CXCL13, CD39, LAYN, CTLA4, RBPJ, DUSP4, GZMA, HAVCR2. |
| Naive-Like T cells (Naïve-like) | CD3D, CD3E, SELL, CCR7, IL7R, TPT1, RPS12, GPR183, RPS18, KLF2, LTB. |
| CD4 Regulatory T cells (Treg) | CD3D, CD3E, CD4, FOXP3, TNFRSF4, IL1R1, IL2RA, CCR6, TNFRSF18, BATF, CTLA4, CCL20, DNPH1, LTB, IL32. |
| CD4 Effector memory T cells (CD4_Tem) | CD3D, CD3E, CD4, JUNB, GZMK, GPR183, KLRB1, CTLA4, ICOS, STAT3, CD28, PDCD1. |
| Mucosal associated invariant T cells (MAIT) | CD3D, CD3E, CD4, CD40LG, IL7R, LTB, KLRB1. |
| Natural killer T cells (NK) | TYROBP, FCER1G, KLRF1, FGFBP2, FCGR3A, GNLY, KLRD1, TRDC, PRF1, NKG7, GZMB, IFNG. |
| Gamma delta T cells (Tgd) | CD3D, CD3E, TRDC, TRGC1. |

**Table Supplementary 1. Gene signatures used to annotate T cell subtypes.**
